# Purified Zymogens Reveal Mechanisms of Snake Venom Metalloproteinase Auto-Activation

**DOI:** 10.1101/2025.06.27.661748

**Authors:** Sophie Hall, Iara Aimê Cardoso, Mark C. Wilkinson, Maria Molina Carretero, Srikanth Lingappa, Bronwyn Rand, Dakang Shen, Johara Boldrini-França, Richard Stenner, Stefanie K. Menzies, Georgia Balchin, Konrad Kamil Hus, Renaud Vincentelli, Andrew Mumford, Alastair W. Poole, Nicholas R. Casewell, Imre Berger, Christiane Schaffitzel

## Abstract

Snake venoms contain diverse mixtures of toxins that evolved to incapacitate prey, but in humans they cause extensive pathology following snakebite envenomation. In viper venom, some of the most potent toxins are the haemorrhagic and coagulopathic snake venom metalloproteinases (SVMPs). Because venoms contain a SVMP cocktail, and due to their cytotoxicity, SVMP characterizations have been hampered by the lack of purified enzymes. By incorporating their prodomain, which blocks the active SVMP site, we overcame their cytotoxicity and enabled recombinant production of zymogens from all three structurally variable SVMP classes (PI, PII and PIII) using our baculovirus/insect cell expression system. Zymogens were auto-activated by incubation with Zn^2+^ ions, resulting in prodomain cleavage, PII disintegrin cleavage and PIII prodomain proteolysis. Auto-activated SVMPs were characterized using protein substrate degradation, platelet aggregation and blood coagulation assays, benchmarked to native venom-purified SVMP. Our recombinant zymogen production protocol is generically applicable for the expression of SVMPs, unlocking biomedical use in haematology, and discovery of novel snakebite therapeutics.

## INTRODUCTION

Snake venom is a complex cocktail of more than 100 proteins and peptides belonging to various toxin families (*1*). Although the number and abundance of isoforms within these families vary extensively across different venomous snakes, the snake venom metalloproteinases (SVMP) are a pathologically important toxin family present in almost all snake venoms. SVMPs play a significant role in the physiological effects resulting from both local and systemic envenoming, such as haemorrhage, coagulopathy and tissue necrosis (*2*). In many medically important viperid snake genera such as *Echis* (saw-scaled vipers), SVMP toxins are the most abundant toxin constituents, on average accounting for 33% of all toxins, though this can be as high as 72% in *Echis ocellatus* (*3–5*) (Fig. 1a).

**Figure 1:**
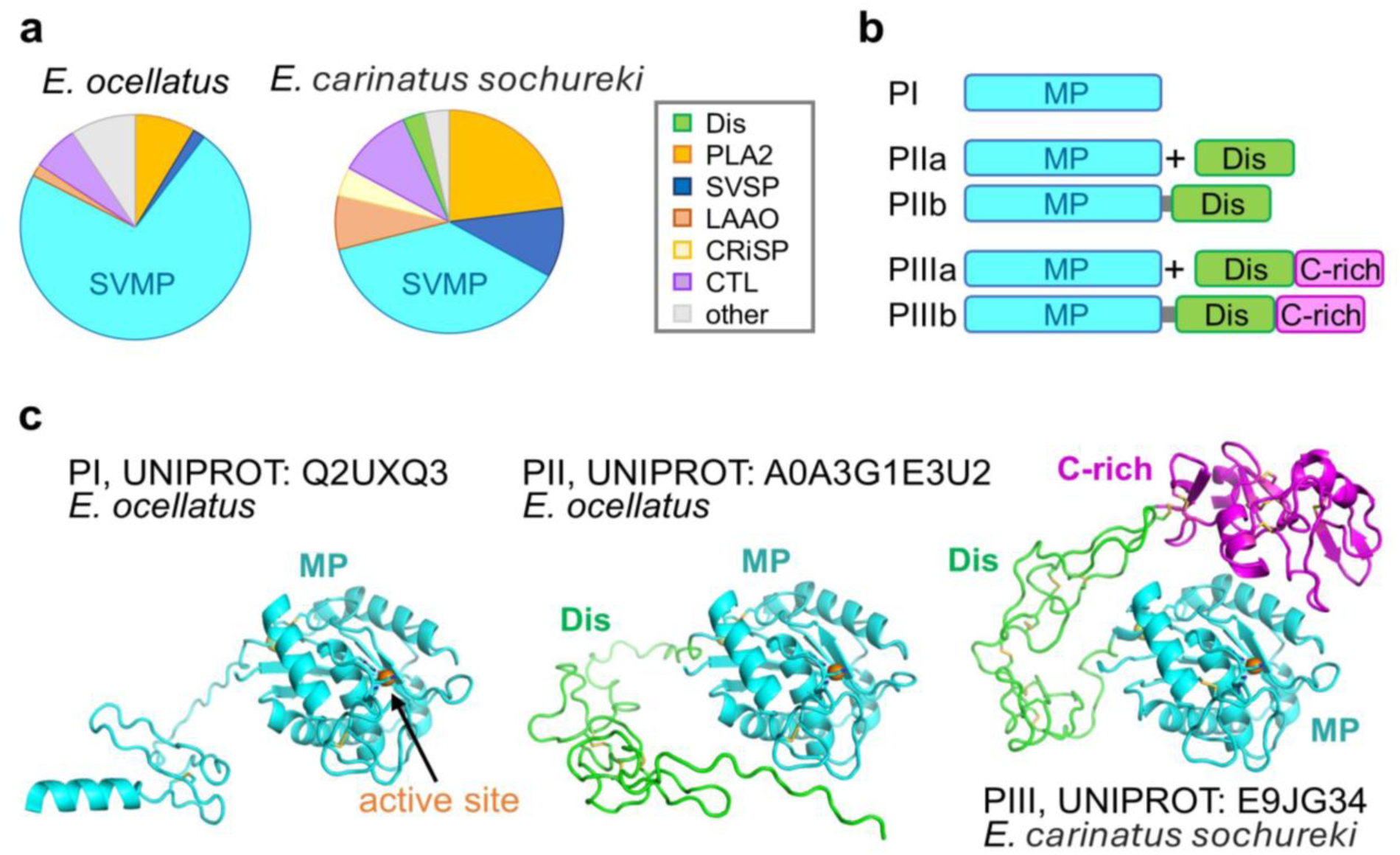
SVMP classification and abundance in *Echis* venom. (**a**) Pie charts displaying *Echis ocellatus* and *Echis carinatus sochureki* venom composition, with SVMP coloured in cyan (*1*). (**b**) Simplified schematic of mature PI, PII and PIII SVMP architecture, showing metalloproteinase domain (MP, cyan), disintegrin domain (Dis, green) and cysteine-rich domain (C-rich, magenta). (**c**) AlphaFold 3 predicted structural models of PI and PII SVMPs from *Echis ocellatus*, and PIII SVMP from *Echis carinatus sochureki*. Domain names and colour coding as in panel b. Zn^2+^ ions (orange) in the active site are shown as spheres.

SVMPs are complex, structurally variable, cysteine-rich, Zn^2+^-dependent metalloproteinases that can be classified into three sub-classes based on domain architecture: PI (comprising metalloproteinase domain only), PII (metalloproteinase and disintegrin domains) and PIII (metalloproteinase, disintegrin-like and cysteine-rich domains) (Fig. 1b); representatives of all three sub-classes are present in different viper venoms (*6*) (*7*).

SVMPs are particularly toxic proteins with a wide range of substrates. They are known to degrade the basement membrane and extracellular matrix proteins, as well as several blood clotting factors, most notably fibrinogen, through their metalloproteinase domain (*8, 9*). Extracellular matrix breakdown and loss in blood capillary wall integrity results in vascular leakage, resulting in both local and systemic haemorrhage (*10*) and in the spread of other venom toxins (*11*). In PII SVMPs, the disintegrin domain contains a charged, integrin-binding motif, most commonly an arginine-glycine-aspartate (RGD) motif, which binds, among others, the fibrinogen receptor integrin α_IIb_β_3_ (*12*), causing inhibition of platelet aggregation.

Because viper SVMPs often target components of the coagulation cascade with unique specificity and/or lack of reliance on typically essential co-factors, these enzymes are highly valuable for investigating and monitoring the treatment of pathologies related to haemostasis (e.g. factor deficiencies, bleeding disorders) (*13*). In the clinic, the SVMPs RVV-X and Ecarin are both used as standards for coagulation tests (*14*). In addition, SVMP-derived disintegrins have been repurposed for use as, or inspiration for the design of, anti-platelet therapies to treat atherothrombosis in the form of unstable angina and myocardial infarction (*1*). Thus, a single snake venom toxin family has yielded several translationally valuable tools for diagnosis and treatment of human disease. However, further exploitation of SVMPs is hampered by the considerable challenge in purifying individual SVMP proteins from venom, and the lack of robust, scalable, recombinant toxin production protocols, representing a bottleneck.

Currently, SVMPs are typically purified from venom by size exclusion chromatography (SEC), followed by ion exchange chromatography of the peaks containing SVMPs. Further separation can be achieved by hydrophobic interaction chromatography or reversed-phase high performance liquid chromatography (RP-HPLC) (*15, 16*). However, complete separation of endogenous functionally active SVMPs is difficult to achieve, because: (i) many venoms contain multiple related SVMP isoforms that share similar physicochemical properties, (ii) the resulting yields of each particular SVMP isoform can be very low, and (iii) the use of some desirable separation approaches (e.g. RP-HPLC) can result in SVMP denaturation. Importantly, this procedure is impeded by the limited access to sufficient amounts of extracted snake venom.

Recombinant expression of SVMPs could resolve the bottleneck, but is severely hampered by their cytotoxicity, domain complexity and high number of cysteine residues that complicate protein folding - for example, the PIII SVMPs contain between 20 and 40 cysteines forming disulfides (*17*). Currently, protocols are available for the production of only a select few specific SVMPs (including albocollagenase, bothropasin and Ecarin) (*18–20*). However, these suffer from limitations such as low yield and inactive proteins. Zymogen production of SVMPs has been described before, but it remains generally elusive how the zymogen is processed into the mature, active SVMP. Current protocols describe incubation of the zymogen for >7 days (*20*), incubation with other active proteinase toxins (*21*), or the insertion of a TEV protease cleavage site into the SVMP amino acid sequence (*22*).

Here, we establish a novel and generic protocol to access functional PI, PII and PIII SVMPs in the quality and quantity required for characterizations and show that all three classes of SVMPs can auto-activate in the presence of zinc ions, resulting in distinct outcomes.

We used our MultiBac baculovirus/insect cell expression system (*23*) for recombinant SVMP zymogen production. The MultiBac system is well-established for secretion of proteins, including the C-terminal heavy chain domain of the highly toxic protein, clostridial botulinum neurotoxin (*24*). We postulate that this system is highly suitable for SVMP production because (i) folding of the multi-domain SVMP zymogen will be supported by eukaryotic, endogenous chaperones present in insect cells and (ii) secreted SVMPs will undergo cellular protein quality control in the endoplasmic reticulum (ER) and Golgi apparatus for correct disulfide bond formation and glycosylation before secretion, ensuring that the toxins produced are correctly folded, soluble and authentically modified. We show that zymogen expression overcomes SVMP cytotoxicity, and we demonstrate successful production of PI and PII SVMPs from *Echis ocellatus* and a PIII SVMP from *Echis carinatus sochureki* (Fig. 1b,c). We establish auto-activation of the SVMP zymogens and concomitant cleavage of the prodomain (all classes), C-terminal tags (PI, PII) and the disintegrin domain (PII). The PIII prodomain is completely degraded by the activated SVMP. We further analyse the cytotoxicity and haemotoxicity of the auto-activated SVMPs *in vitro* using established casein (*25*), fibrinogen (*25*), fluorogenic peptide (*26*) and insulin degradation (*15*) assays. For the PI SVMP, we show that the substrate specificity of recombinant SVMP is identical to that of the corresponding toxin purified from venom, compellingly validating the utility of our strategy.

Our generic SVMP production protocol thus paves the way to systematically characterize SVMPs and unlock biomedical applications, in haematology, thrombosis and inflammation research and as targets for the discovery of novel snakebite therapeutics.

## RESULTS

### Recombinant SVMP zymogen expression overcomes their cytotoxicity

For our expression trials, we chose PI, PII and PIII SVMPs which, according to transcriptomics (*27*), are highly expressed in the venom gland of medically important saw-scaled vipers. We generated baculovirus/insect cell expression constructs for Q2UXQ3 (PI), A0A3G1E3U2 (PII) and E9JG34 (PIII) (Fig. 1c), each encoding a signal sequence for secretion, the toxin of interest and C-terminal tandem octa-histidine- and Avi-tags, codon optimized for both *Spodoptera frugiperda* and *Trichoplusia Ni* as expression hosts (Fig. S1). Initially, recombinant production of mature SVMPs, without the N-terminally fused prodomain, was attempted (Fig. S2a). Protein expression yields, however, were extremely low, primarily due to SVMP cytotoxicity causing pervasive cell death immediately upon baculoviral infection due to initial trace amounts of SVMP. Subsequent expression strategies to produce SVMPs were aimed at inactivating the metalloproteinase by adding an inhibitor, native propeptides or SVMP active site mutants (Supplementary text). Marimastat, a broad-spectrum metalloproteinase inhibitor (*28*) could not prevent cell death induced by PIII SVMP expression. On the contrary, the cell viability in the presence of 3 µM and 6 µM Marimastat was even further decreased (Fig. S2b). Similarly, active site mutants in the PIII SVMP still showed significant toxicity, markedly reducing yellow fluorescent protein (YFP) levels, which serve as a proxy for heterologous protein production, as well as cell viability (Fig. S2c).

In the snake venom gland, SVMPs are produced with an N-terminally fused prodomain that is removed through cleavage during post-translational processing of the protein (Fig. 2a). The prodomain comprises a small, highly conserved propeptide sequence (usually PKMCGVT) - this propeptide acts as a cysteine switch to maintain enzyme latency (*29*). AlphaFold 3 (*30*) predicts that the prodomain adopts a β-barrel structure and folds around the metalloproteinase (MP) domain, slotting the propeptide into the active site of the MP domain, where the propeptide cysteine co-ordinates the active site Zn^2+^ ion, along with the three active site histidines (Fig. 2a). This renders the MP inactive as the Zn^2+^ ion cannot be accessed by substrates in the active site.

**Figure 2:**
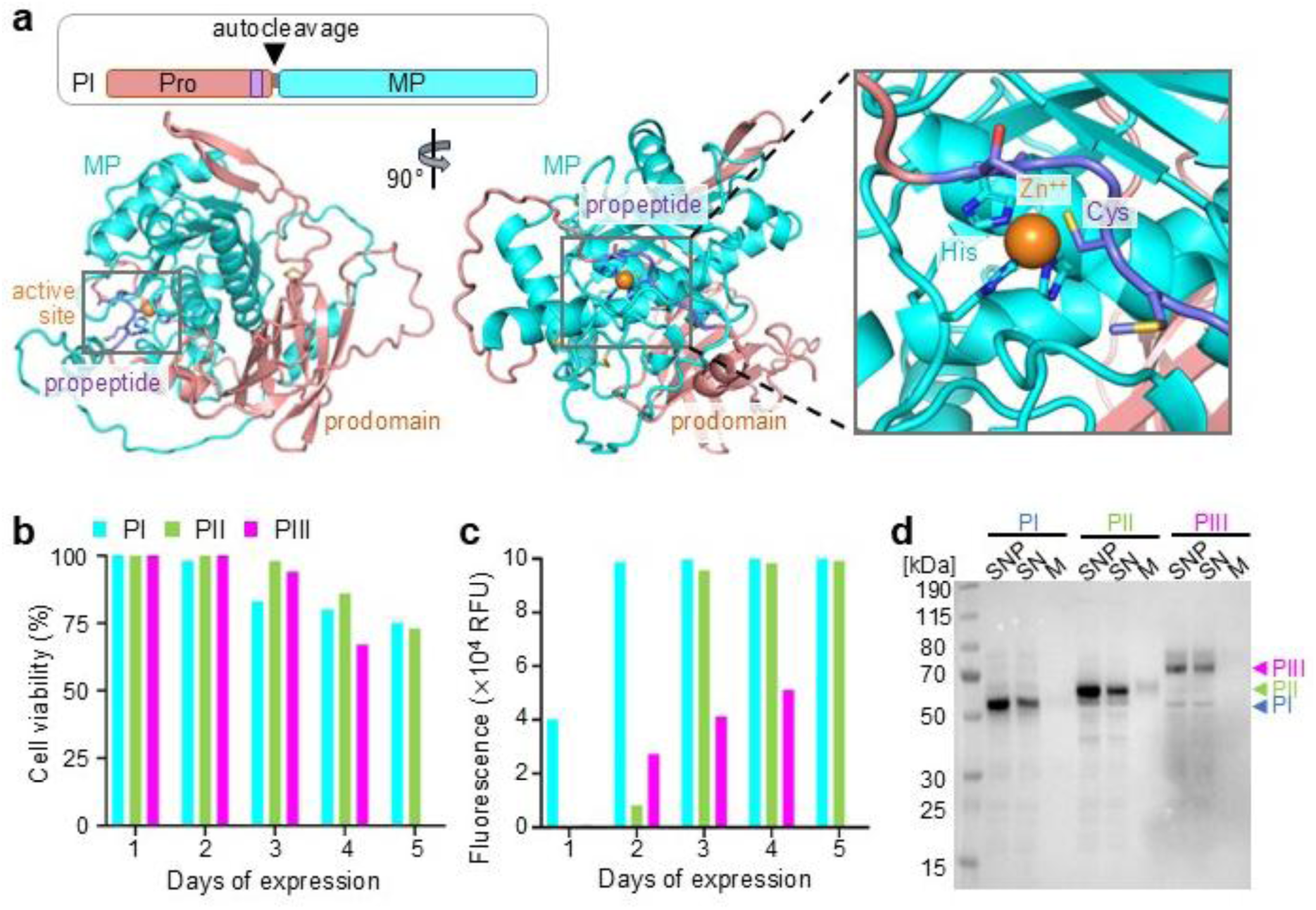
PI, PII and PIII SVMP zymogen expression. (**a**) Above: Schematic of PI SVMP zymogen with N-terminal prodomain (Pro, salmon), propeptide (purple). Below: left: two views of AlphaFold 3 prediction of PI zymogen model; right: zoom in on active site Zn^2+^ ion coordinated by histidines and propeptide cysteine. Expression of SVMP zymogens monitored by (**b**) cell viability, (**c**) YFP fluorescence (PIII harvested on day 4), (**d**) Western blot analysis using HRP-conjugated anti-Penta-His antibody. SNP: supernatant+pellet, SN: supernatant, M: media (10x concentrated). Grey bars: non-toxic protein expression control, cyan: PI SVMP, green: PII SVMP, magenta: PIII SVMP. SVMP zymogen expression was repeated five times.

Inspired by this, we designed propeptide-SVMP fusion proteins with three propeptide repeats to the N-terminus of the SVMP followed by a TEV protease cleavage site. We hypothesized that this would increase the avidity of the propeptide to the MP active site and efficiently inactivate the enzyme. While cell viability and YFP fluorescence levels for PI and PIII slightly increased (Fig. S2d), compared to expression of the mature SVMP (Fig. S2a), YFP expression was still significantly lower as compared to a positive control which expressed an unrelated, non-toxic protein. Western blot analysis confirmed soluble expression of PI and PIII SVMPs (Fig. S2d), but protein yields in the media remained too low to progress to purification.

Therefore, we produced complete SVMP zymogens, where the native signal sequence and the entire prodomain including propeptide is expressed N-terminally to the MP domain (Fig. 2, Fig. S1), adopting a compact structure as predicted by AlphaFold 3 (*30*) (Fig. S3). Cells expressing PI, PII and PIII SVMPs as zymogens showed clearly improved cell viability, remaining above 60% on day 4 of expression (Fig. 2b). PIII SVMP zymogen was harvested on day 4 to avoid additional cell death and lysis. In contrast, the cells expressing PI and PII SVMPs remained ∼75% viable on day 5 of expression when the SVMPs were harvested. Notably, the zymogen-expressing cells had considerably higher YFP readings as compared to the cells previously expressing mature PI and PIII SVMPs, and no loss of YFP fluorescence over time was observed (Fig. 2c). Western blot analysis using an anti-His-tag antibody confirmed the presence of PI, PII and PIII zymogens in cell extract and lysate; and zymogen protein was also detected in the concentrated media fraction, indicating successful secretion (Fig. 2d). We conclude that SVMPs require the full-length N-terminal prodomain to ensure enzyme latency is maintained, and to allow for significant amounts of protein to be produced and secreted when recombinantly expressed using a baculovirus/insect cell system.

### SVMP zymogens can be produced in milligram amounts

Following secretion of the SVMP zymogens into the media, a three-step purification was performed (Fig. S1). Firstly, cells were pelleted by centrifugation, and the media containing secreted proteins were passed through a HiTrap IMAC column for metal affinity purification to enrich the secreted SVMPs from the comparatively large culture media volume. SVMPs were eluted using imidazole. Fractions containing SVMP zymogen were dialyzed and further purified by anion exchange chromatography (IEX), followed by size exclusion chromatography (SEC). SVMP zymogens generally eluted in one prominent peak in SEC (Fig. 3).

**Figure 3:**
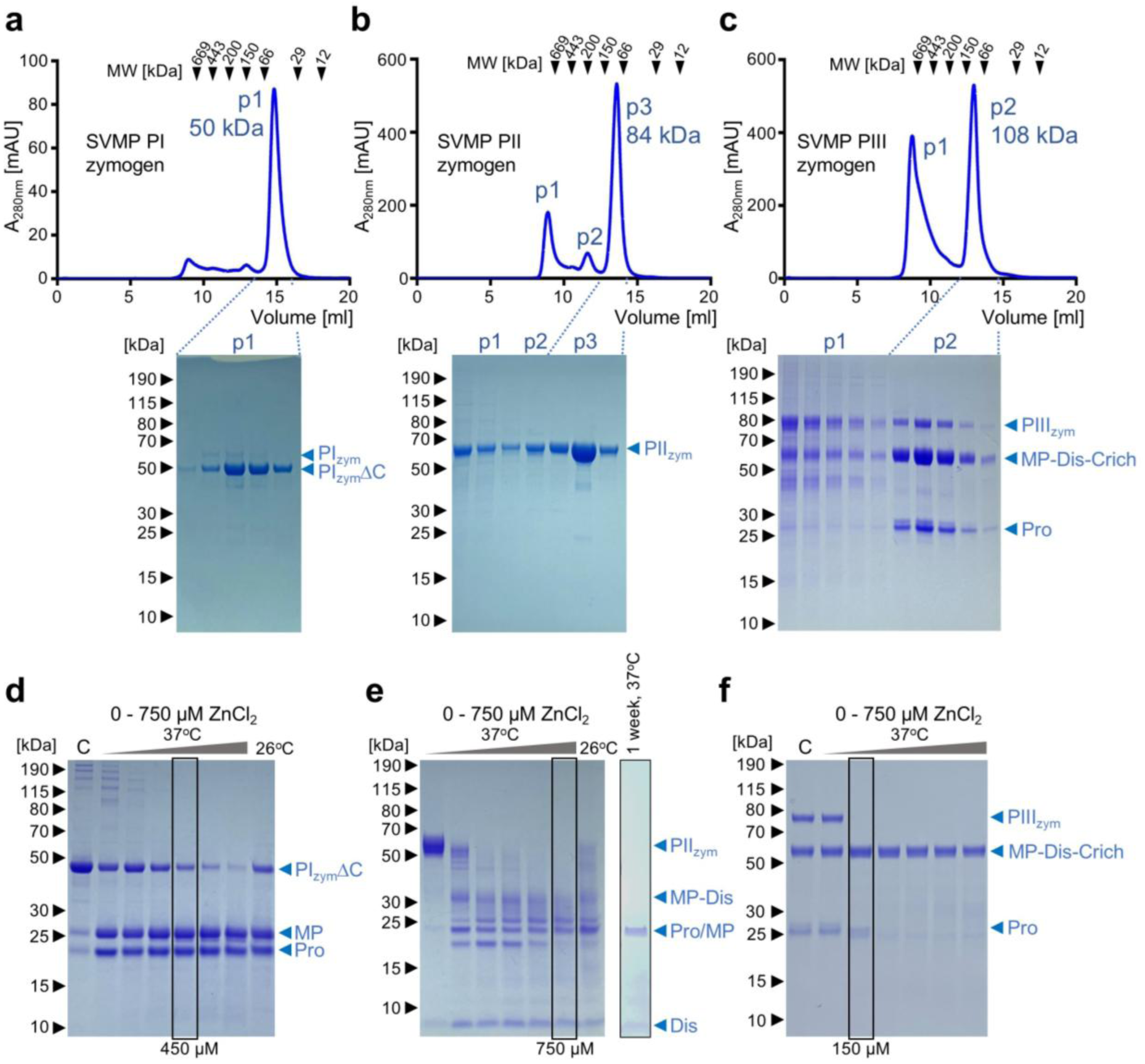
SEC purification of SVMP zymogens and auto-activation. Size exclusion chromatograms and SDS-PAGE of (**a**) PI zymogen (PI_zym_ΔC lacks C-terminal tags), (**b**) PII zymogen, and (**c**) PIII zymogen after IMAC and IEX. PIII_zym_ partial cleavage separates the prodomain (Pro) and the mature PIII. Elution volumes of MW calibration markers are indicated in the chromatograms as black arrows. (**d**) Activation of PI zymogen into metalloproteinase and prodomain. C (control): no 18-hour incubation. (**e**) Activation of PII zymogen into metalloproteinase-disintegrin and disintegrin domain (prodomain cannot be definitively identified). After 1 week only 8 kDa and 23 kDa bands remain. (**f**) Activation of PIII zymogen into prodomain and mature PIII. At higher Zn^2+^ concentrations, the prodomain is degraded. C (control): no 18-hour incubation. SVMP zymogen purifications and activations were repeated five times.

Notably, dialysis following IMAC affinity purification of PI SVMP resulted in self-cleavage of the zymogen as evidenced by a downward shift of the PI protein band in the SDS-PAGE gel from 55 kDa to 45 kDa before and after dialysis, respectively (Fig. S4a). Western Blot analysis indicated a loss of the His-tag after dialysis (Fig. S4b). The protein was confirmed to be PI SVMP by mass spectrometry (Fig. S4c,d). The observed auto-activation of PI SVMP is likely due to residual metal ions in the insect cell media and buffers, trace metal ions leaching off the resin during IMAC purification, and/or a change in pH from the cell culture medium at pH 6 to the purification buffer at pH 8, which is the pH recommended for binding to the IMAC column - the lower pH of the medium being similar to that of the venom gland (pH 5.4), which helps maintain enzyme latency (*31*). The PI SVMP zymogen without the C-terminus (including the Avi-tag and His-tag), eluted from the SEC column in one peak at 14.8 ml, corresponding to a calculated molecular weight (MW) of ∼50 kDa which agrees with the predicted MW for the monomeric zymogen (Fig. 3a). The yield of purified PI SVMP zymogen was 0.63 mg from a 1 L Hi5 insect cell culture.

Following observation of the loss of the SVMP PI C-terminus during purification, a new protein construct was designed with the Avi- and His-tags fused C-terminally to the observed proteolytic degradation site as suggested by MW loss seen by SDS-PAGE and based on AlphaFold 3 models (*30*), giving rise to a shorter construct named PIΔC. In fact, this new C-terminus of the MP domain aligns with that of fully characterized PI SVMPs found in venom (*32–34*). During PIΔC IMAC affinity purification, the zymogen (predicted MW: 45.4 kDa) underwent partial activation, resulting in cleavage between the MP (theoretical MW: 26.2 kDa) and prodomain (theoretical MW: 19.3 kDa) (Fig. S5a). Subsequent IEX allowed for the isolation of a small amount of PIΔC MP domain alone, however the bulk of the PIΔC MP remained bound to the prodomain, forming a stable complex despite cleavage. The PIΔC MP domain alone eluted from the SEC column at a MW of ∼29 kDa, corresponding to the MW expected for the monomeric MP domain (Fig. S6a). The PIΔC metalloproteinase in complex with the prodomain eluted from the SEC column in a different peak corresponding to a MW of ∼48 kDa, confirming that the cleaved prodomain and MP domain form a stable assembly with a 1:1 stoichiometry (Fig. S6b). The yield of PIΔC was 0.44 mg for the MP domain and 3.25 mg for the MP with non-covalently bound prodomain from a 1 L Hi5 insect cell culture.

The PII SVMP zymogen was purified as intact zymogen and did not undergo any (self-) cleavage throughout purification (Fig. 3b). In SEC, a small amount of protein eluted in the void volume, likely due to the very high concentration of PII zymogen when loaded onto the column, possibly causing some aggregation. The majority of PII SVMP zymogen eluted with a small peak corresponding to ∼198 kDa (p2) and a large, sharp peak corresponding to ∼84 kDa (p3). SDS-PAGE analysis with non-reducing loading dye suggests that peak p2 corresponds to a PII zymogen dimer (Fig. S7a,b). Peak p3 corresponds to the monomeric PII SVMP zymogen, which has a predicted MW of 56 kDa, in agreement with its elution volume in SEC. The yield of PII SVMP zymogen was 8.3 mg (peak 3) from a 1 L Hi5 insect cell culture.

The PIII SVMP zymogen underwent partial activation during the purification process, similar to that of PIΔC SVMP. This was observed following elution from the IMAC column (Fig. S5b) and resulted in the presence of three bands in SDS-PAGE (Fig. 3c), corresponding to the PIII zymogen (predicted MW 70.2 kDa), mature PIII SVMP (MP domain, disintegrin-like domain and cysteine-rich domain) with a predicted MW of 50.6 kDa, and the prodomain with a MW of 19.6 kDa. Interestingly, despite the processing of some PIII molecules, the protein still eluted in one peak from the SEC, at a MW of ∼108 kDa based on the elution volume. This indicates that the prodomain again remains associated with the mature SVMP following cleavage, as previously observed for the PIΔC SVMP (Fig. S5a). The yield of the PIII SVMP was 2.2 mg from a 1 L Hi5 insect cell culture.

We observed a difference in the calculated MW based on SEC elution volume (∼108 kDa) and the predicted MW (70 kDa) for the PIII SVMP zymogen. Similarly, the PIII zymogen and mature PIII SVMP both run at a higher-than-expected MW on SDS-PAGE (∼80 kDa and ∼60 kDa, respectively). In order to explain this difference in apparent MW of the PIII SVMP, we analysed the proteins for the presence of N-glycans. The NetNGlyc 1.0 server predicts 2 glycosylation sites in the mature PIII SVMP (*35*), and we tested all purified SVMP zymogens for glycosylation using Peptide:N-glycosidase F (PNGase F) (Fig. S8). The bands corresponding to PI and PII SVMPs did not change in the presence of PNGase F. However, we observed a downward shift in the MW of the bands corresponding to the PIII zymogen and mature PIII SVMP protein in SDS-PAGE following treatment with PNGase F. De-glycosylation of PIII proteins produced bands at ∼77 kDa and ∼57 kDa, closer to their predicted MWs (Fig. S8c). We thus confirmed glycosylation of our recombinant PIII SVMP, accounting for the observed higher MW. Due to unavailability of the corresponding native PIII SVMP from *E. carinatus sochureki*, a more detailed comparative analysis of glycosylation and comparison of glycosylation patterns between native and recombinant PIII SVMP is precluded.

In conclusion, by using the protocol we have developed, we can produce highly purified recombinant SVMP zymogens, each in mg quantities per liter expression culture.

### Co-expression of protein disulfide isomerases does not improve SVMP zymogen yields or folding

Transcriptomic studies have shown that snake venom glands overexpress several chaperones alongside venom toxins, in particular protein disulfide isomerases (PDI), heat shock proteins and calreticulin (*36*). These chaperones are expected to be essential for the correct folding of complex, disulfide-rich toxins. In particular, PDIs assist in the formation, breakage and rearrangement of disulfide bonds, stabilizing the proteins. Successful folding of venom peptides aided by PDI was previously demonstrated for marine cone snails (*37*). Furthermore, overexpression of PDI during recombinant protein expression has been shown to increase overall protein yield (*38*). Therefore, co-expression of SVMPs with PDI could increase SVMP protein yields enabling more protein to be correctly folded and secreted, potentially reducing protein aggregation. To test our hypothesis, we co-expressed the PII and PIII SVMP zymogens with two PDIs: a human PDI (UniProt ID: P07237) and a snake PDI (from *Echis ocellatus*) (*27*), the latter is overexpressed in venom gland tissue (Fig. S9a). Contrary to our expectations, PDI overexpression in insect cells did not noticeably enhance the expression or folding of our SVMP zymogens, and we did not observe increased SVMP yield or activity (Fig. S9 and Supplementary text).

### Auto-activation of SVMP zymogens reveals cleavage of the prodomains, the PII disintegrin domain and PIII prodomain degradation

Cleavage of the prodomain in native zymogens is thought to result in the removal of the propeptide from the active site rendering it accessible to substrates, with Zn^2+^ as an essential cofactor. We thus hypothesized that activation of our purified zymogens could be achieved by addition of Zn^2+^ ions. Therefore, we incubated the SVMPs with Zn^2+^ ions at 37°C for 18 hours and optimized the Zn^2+^ concentration for activation for each SVMP (Fig. 3d-f).

The PI SVMP zymogen could be close to completely activated by 750 μM ZnCl_2_ (Fig. 3d). When analysed by SDS-PAGE, the ∼45 kDa band corresponding to the zymogen mostly disappears, and two lower MW bands of ∼23 kDa and ∼26 kDa appear, corresponding to the prodomain and MP domain, respectively. The two proteins could not be separated under the native conditions of SEC, indicating that the prodomain remains stably bound to the MP domain, despite cleavage (as observed for PIΔC, Fig. S6b). Separation of the MP and prodomain has occurred during SDS-PAGE, however, due to the denaturing and thiol-reducing conditions used. AlphaFold 3 generated models (*30*) (Fig. 2a) suggest the prodomain is located on the opposite side of the MP domain, distal from the active site. Thus, the bound prodomain is not expected to interfere with the ability of the metalloproteinase to bind and cleave substrates, once the propeptide has been detached from the active site. It is interesting to note that the C-terminal part of the PI SVMP zymogen was cleaved during buffer exchange by dialysis while the zymogen was still intact (Fig. S3a) and assumed to be inactive. The observed C-terminal cleavage could be due to a few auto-activated PI SVMPs or due to contaminating proteases from insect cells still present in the affinity purified protein fractions.

The truncated PIΔC SVMP construct was almost completely cleaved into mature protein during purification, however this protein was not fully active in our assays. For full activity, the PIΔC SVMP was incubated with 450 μM ZnCl_2_ leading to significantly higher activity in casein assays (Fig. S6c). Despite complete cleavage of the zymogen, the resulting prodomain and MP domain could not be separated, as was the case for the full-length PI, and, thus, no MP was isolated from this construct either (Fig. S10). Because this PIΔC SVMP corresponds better to the processed, native version of PI SVMP protein, PIΔC was used in all further activity assays and is referred to as PI SVMP from here on.

The PII SVMP zymogen was completely activated by 750 μM ZnCl_2_ (Fig. 3e). Notably, when analysed by SDS-PAGE, a clear separation of prodomain and mature protein was not observed. The band migrating at ∼40 kDa corresponds to MP and disintegrin (confirmed by Western blot, Fig. S7c). This band disappears with higher concentrations of Zn^2+^ and/or longer incubation times. The band migrating at ∼8 kDa is thought to correspond to the processed disintegrin domain but could not be confirmed by Western blot due to loss of C-terminal tags during activation. After incubation for 1 week, only the 8 kDa and 23 kDa bands remained (Fig. 3e). Based on apparent MW, these bands correspond to the disintegrin and prodomain, respectively. In summary, PII SVMP has undergone further processing to remove the tags and disintegrin from the MP domain, and the MP domain appears to be unstable over time.

The PIII SVMP zymogen was completely activated by the addition of 150 μM ZnCl_2_ (Fig. 3f) as evidenced by the disappearance on SDS-PAGE of the ∼77 kDa band corresponding to the zymogen. Instead, the band strengthened at ∼60 kDa corresponding to the mature SVMP (MP domain plus disintegrin-like domain and cysteine-rich domain) together with a band at ∼25 kDa corresponding to the prodomain. Notably, at higher concentrations of Zn^2+^, the C-terminal tags and the prodomain band are degraded by the PIII SVMP.

We conclude that all SVMP zymogens studied can auto-activate by incubation with Zn^2+^, leading to cleavage of the prodomain from the zymogen. In the case of PIΔC and PIII SVMPs the prodomain remains associated with the SVMP at low Zn^2+^ concentrations. After Zn^2+^ activation, the C-terminal tags and the PIII prodomain are degraded by the activated SVMPs, and in the case of PII SVMP, further proteolysis occurs into the constituent parts.

### Casein and fibrinogen degradation assays demonstrate SVMP activity

Degradation of casein is a general protease activity assay that is used in functional assays for all SVMP classes (*25*). SVMP zymogens were supplemented with optimal concentrations of ZnCl_2_ for SVMP activation (Fig. 3d-f) and incubated with 0.525 μg/μl (∼23 μM) casein overnight at 37°C. The degradation of casein was analysed by SDS-PAGE; indicated concentrations relate to input SVMP zymogen concentrations (Fig. 4a-f). The PI SVMP degraded casein (Fig. 4a); activity was observed when adding 73 nM PI and complete degradation of casein occurred at the addition of 1.17 μM PI. The PII SVMP exhibited weak activity against casein (Fig. 4b), which may be due to the self-degradation of the MP domain observed during Zn^2+^ activation (Fig. 3e). Casein degradation was observed when adding 1.36 μM PII, as evidenced by the emergence of lower MW bands (Fig. 4b). The PIII SVMP was active against all casein chains (Fig. 4c), with activity seen from concentrations as low as 21 nM PIII, and complete digestion of casein was observed with 343 nM PIII.

**Figure 4:**
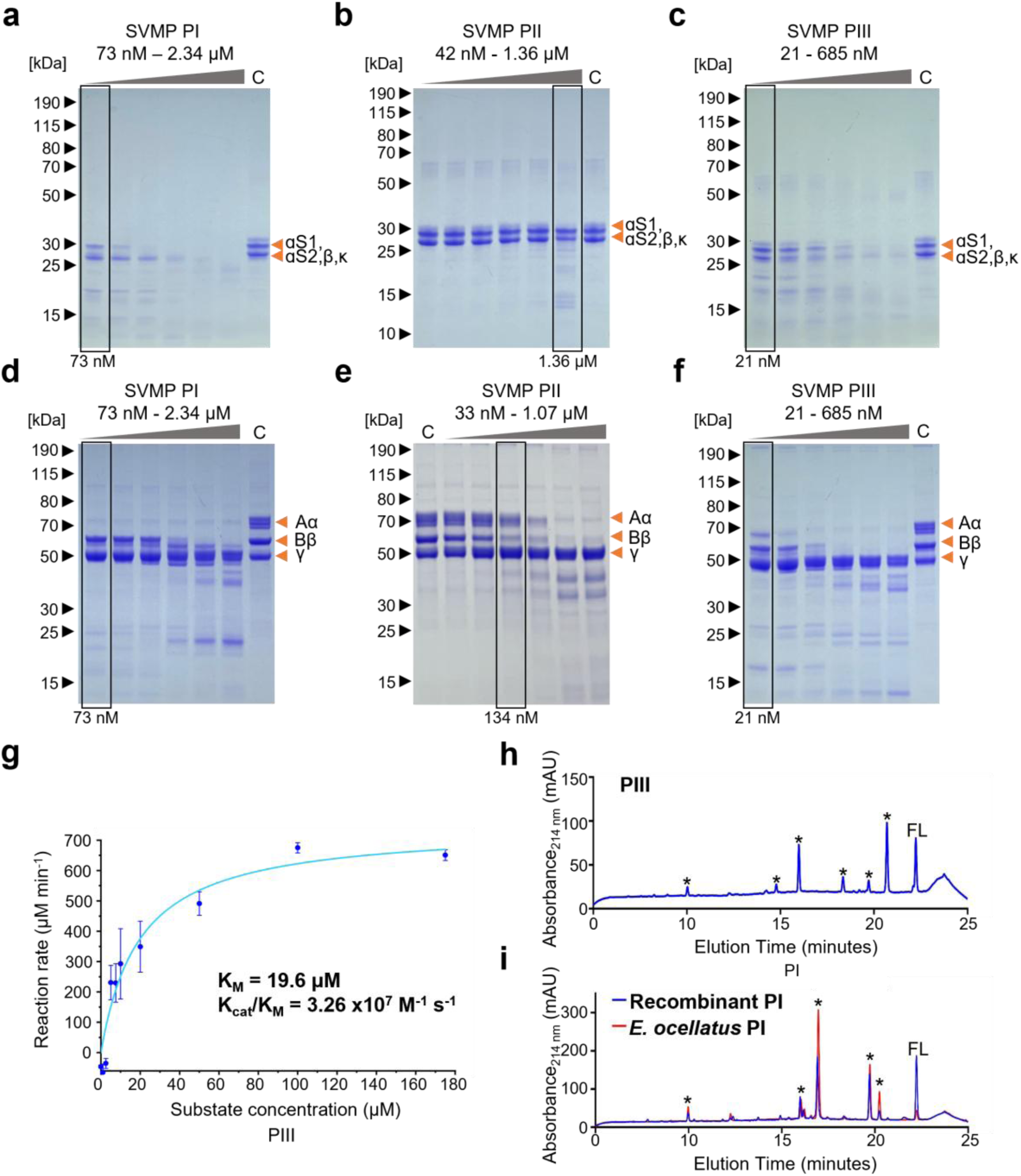
*In vitro* SVMP activity and substrate specificity. Casein degradation in the presence of increasing amounts of (**a**) PI, (**b**) PII and (**c**) PIII SVMPs. Orange arrowheads: domains αS1, αS2, β and κ. Fibrinogen degradation in the presence of increasing amounts of (**d**) PI, (**e**) PII and (**f**) PIII. Orange arrowheads: fibrinogen chains Aα, Bβ and γ. C (control): incubation in the presence of 5 mM EDTA. (**g**) PIII SVMP (20 nM) activity towards the fluorogenic peptide substrate ES010. (**h,i**) Degradation of insulin B by (**h**) PIII and (**i**) recombinant (blue) and *Echis ocellatus* PI (red). Full-length insulin (FL) and degradation products (*) are marked above the peaks. All degradation assays were repeated three times. The fluorogenic peptide assay shows the result of three independent repeats.

Next, we performed fibrinogen degradation assays followed by SDS-PAGE analyses (Fig. 4d-f). None of our recombinant SVMPs had very strong activity against the γ chain of fibrinogen, as is usually the case with SVMPs (*34*). However, PI, PII and PIII SVMPs degraded Aα and Bβ chains of fibrinogen to different extents.

The majority of PI SVMP activity was directed against the Aα chain of fibrinogen (Fig. 4d), with the lowest amount of enzyme tested (73 nM). At higher concentrations of PI (585 nM), activity towards the Bβ chain was also observed. Therefore, PI SVMP can be classified as an alpha/beta fibrinogenase, with a preference for the Aα chain. PII SVMP caused removal of fibrinopeptide B (FpB) from the Bβ chain (Fig. 4e) at a concentration of 134 nM. At higher concentrations (268 nM or more), the Aα chain was also degraded. Thus, PII SVMP is an alpha/beta fibrinogenase with a preference for the Bβ chain. PIII SVMP degraded the Aα chain of fibrinogen at the lowest concentration of PIII tested, 21 nM (Fig. 4f). When supplementing 85 nM PIII or more, the band at 60 kDa corresponding to the Bβ chain disappears as well. Thus, we classify PIII SVMP as an alpha/beta fibrinogenase, with a preference for the Aα chain.

To summarize, all recombinant SVMPs (PI, PII, PIII) degraded casein, confirming their activity. They also degraded the alpha and beta chains of fibrinogen in a fibrinogenolytic manner, rather than cleaving fibrinopeptides in a thrombin-like manner.

### Auto-activated PIII SVMP shows high substrate affinity and catalytic efficiency

We next tested if the SVMPs cleave a quenched fluorogenic substrate, ES010 (R & D Systems) which is commonly used to study SVMPs (*39*) and matrix metalloproteinases (*40*) for small molecule therapeutic discovery. PIII SVMP was able to cleave the Gly-Leu bond in ES010 (Fig. 4g). In contrast, SVMPs PI and PII did not effectively cleave this PIII SVMP and MMP-prototypic substrate under the conditions tested (Fig. S11a). For the PIII SVMP, the rate of reaction (monitored by fluorescence increase) was plotted against varying substrate concentrations (ranging from 0 μM to 180 μM), and a Michaelis-Menten curve was calculated (Fig. 4g). The low K_M_ of 19.6 μM indicates a high binding affinity of the PIII SVMP for the ES010 fluorogenic substrate. The high catalytic efficiency (k_cat_/K_M_) of 3.26 x10^7^ M^-1^s^-1^ further suggests that the PIII SVMP is highly effective and specific in processing this substrate. For comparison, a k_cat_/K_M_ value of 4,471 M^-1^s^-1^ was reported for matrix metalloproteinase MMP10 for an ESO10-derived peptide (*41*), and k_cat_/K_M_ values of 30,000 M^-1^s^-1^ for a triple-helical peptide for MMP-2 (*42*). Together, these parameters demonstrate that the recombinant PIII SVMP exhibits both strong substrate affinity and high catalytic efficiency towards this prototypical PIII SVMP substrate.

### Native and recombinant PI SVMP exhibit the same substrate specificity in insulin B degradation assays

Next, we assayed the activity of SVMPs PI, PII and PIII against insulin B, a test substrate used previously in SVMP characterization (*15*). Degradation of insulin B at multiple sites was visualized by RP-HPLC. PI and PIII showed distinct insulin B degradation patterns indicating different sequence specificities and cleavage sites while the PII SVMP did not degrade insulin B (Fig. S11b). PIII SVMP cleaves insulin B into 6 products (Fig. 4h). PI SVMP cleaves insulin B into 5 products (Fig. 4i). While we could not purify the native PII and PIII SVMPs from *Echis* venom for comparative purposes, the native PI SVMP can be readily purified from *Echis ocellatus* venom (*43*). Therefore, we could compare insulin B cleavage by our recombinant PI SVMP with insulin B cleavage by the native source-purified PI enzyme. Identical degradation products of insulin B were observed (Fig. 4i). Thus, our recombinant PI SVMP is active and has the same substrate specificity as the native enzyme from venom, underscoring that our recombinant expression strategy leads to native-like SVMPs, compellingly validating our approach.

### PII zymogen and activated PII SVMP inhibit platelet aggregation

Some haemorrhagic SVMPs can disrupt cell adhesion and prevent haemostasis by interfering with platelet aggregation (*44*). SVMPs with either a disintegrin (PII), or disintegrin-like domain (PIII) can inhibit platelet aggregation through a motif found in these domains. Most commonly, this corresponds to a RGD sequence, which binds integrins. Binding of disintegrin to integrin α_IIb_β_3_ can be measured through inhibition of 2-MeS-ADP-induced platelet aggregation whereby turbidity is determined by plate aggregometry (Fig. 5a). When tested at a concentration of 500 nM SVMP, platelet aggregation was reduced to 12% and 36% by the activated PII SVMP and the zymogen form, respectively, confirming that the disintegrin is active and capable of inhibiting platelet aggregation. Based on our results, the presence of the prodomain in the zymogen has low inhibitory effect on the RGD-containing disintegrin domain, as anticipated. In comparison, the PI and PIII SVMPs did not inhibit 2-MeS-ADP-induced platelet aggregation (Fig. 5a). The PI SVMP lacks the disintegrin domain and instead of an RGD motif, the PIII SVMP has a divergent ‘RSECD’ motif in its disintegrin-like domain, which most likely does not interact with integrin α_IIb_β_3_. We conclude that our recombinant PII SVMP inhibited platelet aggregation, confirming disintegrin activity in the context of both the zymogen and also the activated PII SVMP.

**Figure 5:**
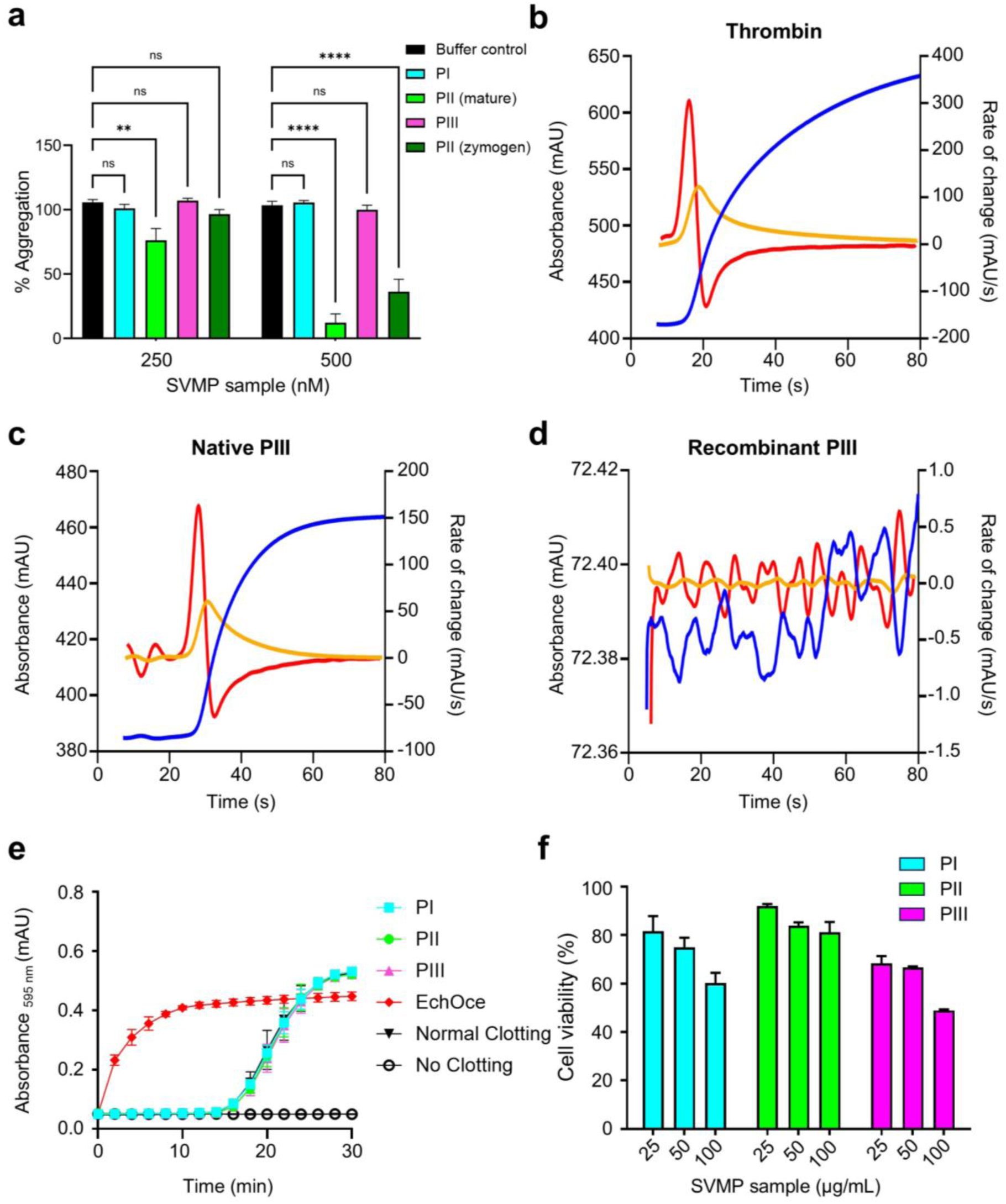
SVMP blood coagulation, platelet aggregation and cytotoxicity. (**a**) Inhibition of platelet aggregation by PI, PII (mature and zymogen) and PIII. Control: PRP with buffer. Data are presented as the mean of % aggregation ± SEM. A two-way ANOVA was conducted, followed by simple effects analysis (Dunnett post-test) to compare column means (SVMPs) within each row (concentrations). ****: *p* ≤ 0.0001, **: *p* ≤ 0.01. The assay was carried out in experimental duplicate and biological triplicate. (**b**)-(**d**) Thrombin clot time assays using the ACL TOP coagulation analyser. Citrated human blood was spiked with (**b**) thrombin, (**c**) a pool of native *Echis ocellatus* PIII SVMPs, and (**d**) the recombinant PIII SVMP. Change in absorbance (blue) is plotted on the left Y axis, rate of change (1^st^ derivative: orange; 2^nd^ derivative: red) is plotted on the right Y axis. Assays were repeated three times. (**e**) Coagulation profiles of the recombinant SVMPs and *Echis ocellatus* venom in bovine plasma over 30 minutes at an absorbance of 595 nm. Data points represent the mean of three individual values ± SD. The experiment was repeated two times. (**f**) MTT cytotoxicity assay showing viability compared to the buffer-only control of HaCaT cells after 24-hour incubation with activated PI, PII and PIII SVMPs. Bars show the results of two independent repeats.

### PIII SVMP has no pro-coagulant activity in thrombin clot time assays

Thrombin clot time assays measure the time required for thrombin to convert fibrinogen to fibrin, leading to clot formation. This assay is used in the clinic to highlight abnormalities in fibrinogen and fibrin clot formation. In our assays, thrombin was replaced with each SVMP to identify the thrombin-like activity of the enzyme. The assay measures turbidity, which is plotted against time. The first and second derivatives are plotted on the same graph, and the maximum rate of change of turbidity (the peak in the 2^nd^ derivative) indicates the time of clot formation. When thrombin is added, fibrin clot formation requires 16.5 seconds (Fig. 5b). As a positive control and example of pro-coagulant SVMPs, thrombin was replaced with native PIII SVMPs purified from *Echis ocellatus* venom. Fibrin clot formation occurs in 28.1 seconds in the presence of the *Echis ocellatus* PIII SVMPs (Fig. 5c), confirming clot formation and pro-coagulation activity. However, when replaced with the recombinant PIII SVMP (from *Echis carinatus sochureki*) no clot formation occurs (Fig. 5d). Similarly, PI and PII SVMPs do not cause blood clotting (Fig. 5e, Fig. S11c). This shows that our recombinantly produced SVMPs have no pro-coagulant activity, which was reported previously for the PI SVMP (*43*) but remained unknown for the PII and PIII SVMPs.

### PI and PIII SVMPs are cytotoxic

HaCaT cells (immortalized human keratinocyte cells), commonly used in dermatological, toxicological and cancer research, were incubated with PI, PII and PIII SVMPs for 24 hours, and cytotoxicity was monitored using the MTT (3-(4,5-dimethylthiazol-2-yl)-2,5-diphenyltetrazolium bromide) assay (*45*). At the tested concentration, PII SVMP showed minimal cytotoxicity (Fig. 5f). In comparison, at 100 µg/mL the PI and PIII SVMPs were clearly cytotoxic, with cell viability decreasing by 40% and 50% after 24 hours, respectively (Fig. 5f). The cytotoxic effect of our PIII SVMP (*Echis carinatus sochureki)* (100 µg/ml) is comparable to that of crude *Echis carinatus* venom on HEK 293T cells, which induces 63% cytotoxicity with 100 µg/ml venom after 3 hours (*46*). Published IC_50_ values for HaCaT cells treated with whole venom for 24 hours fall between 10 and 100 µg/ul (*47*), however it is important to highlight the distinction between the complex cocktail of toxins found in venom and the homogenous, purified SVMP evaluated in this study.

## DISCUSSION

The characterization of SVMP toxins, critical for effective bioprospecting and the discovery of snakebite therapeutics, is severely hampered by difficulty in isolating the native proteins from venom. Here, we established a generic protocol for efficient recombinant production of the three structurally diverse classes of SVMPs as zymogens, which overcomes this obstacle. We demonstrate expression of PI, PII and PIII SVMP zymogens from *Echis* species in high quality and quantity using the MultiBac baculovirus/insect cell expression system.

In SVMP zymogens, the highly conserved propeptide sequence which blocks the active site of the MP domain, is clamped in place by the N-terminal prodomain. It should be noted that despite expression as a zymogen, residual cytotoxicity was still observed, evident by the decrease in cell viability throughout expression and reduction in the amount of co-expressed YFP fluorescence due to cell lysis (Fig. 2). In agreement with this observation, PI and PIII SVMPs were able to partially activate themselves, resulting in the presence of three characteristic bands in Coomassie-stained SDS-gels – the zymogen, mature SVMP and prodomain. We speculate that this auto-cleavage could be due to the SVMP active site accreting divalent metal ions from the media and buffers used. Additionally, it is likely that the change in pH from 6 (expression) to 8 (purification) allows the activation of SVMPs, as is the case when venom is released from the venom gland (pH 5.4) into the victim (pH 7.4) during envenoming (*31*). A complete control of self-activation of SVMP zymogens would be highly desirable. However, previous studies reported that (i) SVMP zymogens are processed within secretory cells of the venom gland (*48*), and that (ii) mature SVMPs accumulate in secretory vesicles during venom production (*49*). Accordingly, complete prevention of SVMP zymogen auto-processing will be very difficult to achieve because this would require Zn^2+^ ion depletion within the cells which would result in cytotoxicity.

Of note, despite the self-cleavage of the PI and PIII zymogens, the majority of the mature SVMP protein remained associated with the prodomain. Interestingly, this did not affect the activity of the SVMPs. According to AlphaFold 3 structure predictions, the prodomain is located on the side of the MP domain, which is opposite to the active site. Upon activation, cleavage of the prodomain releases the propeptide from the active site. This renders the Zn^2+^ ion in the active site free to interact with a substrate. In agreement, we observe that the SVMPs are active even if the prodomain, now no longer covalently linked to the SVMP, remains bound to the MP domain (Fig. 3c, Fig. S5).

SVMP zymogens undergo additional post-translational processing: The PI SVMP construct described here was based on transcriptomic data and corresponds to the UniProt sequence ID: Q2UXQ3. During purification, it underwent C-terminal truncation by ∼10 kDa, leading to a smaller protein which corresponds closely to previously characterized PI SVMPs (*32–34*). Moreover, we observed a general trend of loss of C-terminal tags during purification, activation and subsequent activity assays for all three SVMP classes.

Here, we show that the recombinant PIII SVMP comprises N-linked glycosylation. We would like to point out that when expressed in insect cells, the glycosylation pattern may be different compared to endogenous SVMPs due to differences in the snake and insect glycosylation pathways. The Sf21 and Hi5 cells used in this study typically produce N-glycans that are trimmed to a core ‘paucimannose’ structure (Man3GlcNAc2), often with an alpha1,6-fucosylation (*50*). While snakes can produce more complex, sialylated N-glycans, glycomic studies of native venoms (e.g. *Bothrops* venoms) have demonstrated that high-mannose and paucimannose structures are also prevalent in native SVMPs (*51*). Therefore, the recombinant glycoforms likely represent a subset of the native glycan heterogeneity and not the full diversity found in snake venom toxins. Moreover, Sf21 and Hi5 cells recognize the same N-glycosylation sequon (Asn-X-Ser/Thr) as reptilian cells, and therefore the site-occupancy remains consistent with the native protein, preserving the overall topography of the toxin.

To improve SVMP yields and/or activity, we hypothesized that co-expression of SVMPs with human or snake PDI would aid their proper folding, ensuring the multiple disulfide bonds were correctly formed. Based on transcriptomic data from *Echis* venom gland (*27*), PDIs are highly overexpressed in the snake venom gland. However, PDI over-expression in insect cells did not improve protein yields nor enzyme activity. We speculate that the endogenous insect cell PDIs are sufficient for SVMP folding in our setup. Alternatively, a combination of different chaperones may be required for further boosting SVMP production.

SVMP zymogens from all three classes could be auto-activated by the addition of Zn^2+^ ions. While some PI SVMP was processed during purification, any remaining PI zymogen could be cleaved into the mature SVMP and prodomain by incubation with Zn^2+^. The PII zymogen could be activated by the same means, although there was no clear separation into prodomain and mature protein. Instead, we observed two bands corresponding to the disintegrin and prodomain after prolonged incubation with Zn^2+^ (Fig. 3e). It is unlikely that the observed disappearance of the MP domain is due to misfolding because the PII zymogen is well expressed with high yields (9-10 mg/L culture) in insect cells and eluted in two clean peaks in SEC corresponding to a monomer and dimer, respectively. In the case of misfolding, we would have expected aggregation and/or degradation during purification, and we would not expect to see degradation of the zymogen in SDS-PAGE analysis only after addition of Zn^2+^ which activates the metalloproteinase (Fig. 3e). Notably, the disintegrin domain of this PII has 100% sequence identity to ocellatusin, a short, monomeric disintegrin of *Echis ocellatus* venom (*52*). Previous studies have identified a discrepancy between the amounts of ocellatusin and its PII-derived metalloproteinase in the venom, whereby ocellatusin represents 3.9%, and the SVMP less than 0.1% of venom proteins (*3*). In summary, the disintegrin is likely liberated from this PII SVMP and is the more pathogenic component of the SVMP, causing inhibition of platelet aggregation (Fig. 5a). In contrast, the PII-derived metalloproteinase domain is unstable and therefore requires high toxin concentrations to elicit proteolytic activity in our activity assays (Fig. 4b,e).

The recombinant PIII zymogen was fully auto-activated in our experiments by the addition of Zn^2+^ ions (Fig. 3f). Interestingly, this PIII SVMP was able to degrade its prodomain despite the latter initially remaining bound to the mature protein according to SEC (Fig. 3c). This agrees with the finding that prodomains, which are highly conserved in SVMPs, are rarely identified in the venom proteome (*53*), suggesting that all prodomains might eventually be fully degraded by venom proteases including PIII SVMPs.

Activated recombinant SVMPs were active against bovine casein, a general protease substrate (Fig. 4a-c). PI and PIII exhibited strong activity, degrading all bands, while PII was significantly less active, requiring much higher enzyme concentrations. This agrees with our previous observation that bands corresponding to the MP-disintegrin protein or the MP domain alone after activation with Zn^2+^ (Fig. 3e) are absent in SDS-PAGE, indicating that the MP domain is likely unstable.

Fibrinogen degradation assays are particularly relevant for SVMPs and indicate their potential haemotoxic activity (Fig. 4d-f). The 340 kDa fibrinogen glycoprotein is made up of three pairs of polypeptide chains, which lead to three groups of distinct bands when analysed by reducing SDS-PAGE: the Aα (70-75 kDa), Bβ (65 kDa) and γ (50 kDa) chains. In contrast to the endogenous serine protease thrombin, which cleaves fibrinopeptides A and B off the Aα and Bβ chains, causing self-polymerization into fibrin fibres and formation of blood clots (*54*), the PI, PII, and PIII SVMPs studied here caused a fuller degradation of fibrinogen (Fig. 4d-f). All SVMPs were active against both the Aα and Bβ chains of fibrinogen; the PI and PIII showed a preference for the Aα chain and the PII preferentially degraded the Bβ chain of fibrinogen. We conclude that the three SVMPs interfere with clot formation and would thus contribute to an anticoagulant outcome upon envenoming. In agreement, the PIII SVMP cannot cause clot formation in the thrombin clot time assay (Fig. 5d), suggesting it is not an Ecarin-like prothrombin activator, which is typically responsible for procoagulant activity seen in such assays with *Echis* venoms (Fig. 5c,e).

The recombinant PIII SVMP was highly active against a fluorogenic peptide substrate, ES010, showing high affinity and specificity for cleavage of the Gly-Leu bond within the peptide (Fig. 4g). In this context, we would like to point out that enzyme concentrations and kinetics are based on zymogen concentrations, assuming 100% SVMP activation by Zn^2+^ (Fig. 4a-g). If zymogen activations were incomplete or non-linear, our k_cat_/K_M_ calculations could either underestimate or overestimate the PIII SVMP’s catalytic efficiency (Fig. 4g).

The SVMP PIII was also active against insulin B (Fig. 4h), as was SVMP PI (Fig. 4i) which produced identical degradation products to native PI, confirming identical cleavage sites and successful recombinant production of toxin with native-like characteristics, validating our approach.

In addition to the proteolytic activity of the MP domain, PII and PIII SVMPs also include a disintegrin or disintegrin-like domain containing a potential integrin binding motif. In platelet aggregation assays, our RGD-containing PII SVMP inhibited 2-MeS-ADP-induced platelet-aggregation with and without the prodomain present (Fig. 5a). The RSECD-containing PIII, in contrast, did not inhibit 2-MeS-ADP-induced platelet aggregation, but this does not exclude other disintegrin-like activity of this domain.

Zymogen expression was previously reported for ADAMs (A Disintegrin and Metalloproteinases) and MMPs (Matrix Metalloproteinases), which are structurally and functionally related to SVMPs. ADAMs and MMPs are involved in cell-cell interactions, ectodomain shedding and degradation of extracellular matrix components respectively (*55*). ADAMs and MMPs have become prominent targets in drug discovery (*56, 57*). For MMPs, zymogen expression in mammalian and insect cells has been reported and enzyme activation was achieved via 4-aminophenylmercuric acetate or trypsin to remove the prodomain (*58, 59*). Moreover, ADAMs have been expressed with their prodomain to aid proper folding and latency (*60, 61*). The ADAMs prodomain has been removed post-translationally, e.g. by co-expression of furin (*62*). Interestingly, in agreement with our PI and PIII SVMP observations, ADAM33 zymogen was processed and secreted in *Drosophila* S2 cells, with the prodomain remaining associated with the catalytic domain after purification. The processing was enhanced by addition of cadmium chloride and zinc chloride (*63*). Auto-activation of zymogens by Zn^2+^ incubation alone, however, has not been reported to the best of our knowledge.

While ADAMs and MMPs are well characterized in the literature, comparable insights into SVMPs were largely absent to date, due to lack of means for recombinant SVMP protein production. Our generic protocol for recombinant SVMP zymogen production of all three classes overcomes their high cytotoxicity and allows, for the first time, to study auto-activation revealing important differences between the SVMP classes. Furthermore, our protocol will unlock the production of many additional hitherto inaccessible SVMPs, allowing in-depth characterization of their post-translational processing, enzymatic activity, substrate specificity, cyto- and haemotoxicity, and enable the evaluation of potential biomedical applications to develop new diagnostics and treatments for blood clotting disorders. Our study paves the way to engineer existing and new SVMPs for improved and novel applications in haematology, thrombosis, and inflammation research and other important areas in which SVMPs can play a crucial role. Importantly, the recombinantly produced, tagged SVMPs can serve as antigens for the future selection and evolution of neutralizing antibodies as a basis for next-generation snakebite treatment to combat snakebite envenomation.

## MATERIALS AND METHODS

### SVMP expression construct design

PI, PII and PIII SVMP constructs were designed by fusing an Avi- and His-tag (GGSGLNDIFEAQKIEWHEHHHHHHHH*) to the C terminus of the native SVMP sequences listed on UniProt (RRID: SCR_004426) (PI – Uniprot ID: Q2UXQ3, PII – Uniprot ID: A0A3G1E3U2, PIII – Uniprot ID: E9JG34). The sequences were codon-optimized for *Spodoptera frugiperda*, gene synthesized (Genscript) and inserted into the plasmid pACEBac1 (Geneva Biotech), which acted as the donor in the MultiBac system (*64*). Structure predictions of SVMP zymogens were performed using AlphaFold 3 (*30*) in the presence of a single Zn^2+^. Per residue predicted Local Distance Difference Test (pLDDT) plots, indicating local confidence, were generated from B-factor values stored in the .cif files. The highest-ranked model among the five predictions was selected for visualization and figure generation using UCSF ChimeraX (RRID:SCR_015872, https://www.cgl.ucsf.edu/chimerax/).

Mature SVMP constructs did not include the prodomain, but did contain an N-terminally fused melittin signal sequence (MKFLVNVALVFMVVYISYIYA). Propeptide-SVMP fusions contained an additional N-terminally fused propeptide triplicate and TEV site (GGSPKMCGVTPKMCGVTPKMCGVTENLYFQSN) following the melittin signal sequence. SVMP active site mutants were based on the mature SVMP constructs, but with the active site glutamic acid mutated to aspartic acid and glycine mutated to cysteine (*65*). Zymogen constructs comprised the native signal sequence preceding the native prodomain (which is specific to the SVMP) and the SVMP sequence with a C-terminal Avi-His-tag fusion. PI was found to undergo C-terminal processing, the site of which was predicted by modelling using AlphaFold 3 (*30*). PIΔC construct contained the Avi-His-tag fusion following this site.

### Recombinant SVMP expression

Each SVMP-containing pACEBac1 plasmid was expressed using the MultiBac baculovirus/insect cell expression system following established protocols (*66*). SVMPs were expressed in Hi5 insect cells at 19°C, in ESF 921 Insect Cell Culture Media (Expression Systems). The cells were monitored throughout expression by measuring YFP fluorescence (encoded in baculoviral genome) and cell viability (by determining percentage of alive cells using Trypan Blue). A control expression of an unrelated, non-toxic protein (E3 ligase) was used as a control for YFP and cell viability measurements. Expression in the presence of Marimastat was tested using a final concentration of 0 μM, 3 μM and 6 μM Marimastat in the cell culture media, which was added alongside the virus (at the start of expression). Cultures were harvested 5 days after infection by centrifugation (1000 xg, 10 minutes), or when the cell viability dropped below 80% if this occurred sooner. The secreted SVMP proteins were purified immediately after harvesting from the supernatant.

### Western blotting

Proteins resolved by SDS–PAGE were transferred to PVDF membranes (Trans-Blot Turbo mini 0.2 µm PVDF Transfer Pack, Bio-Rad) using a Trans-Blot Turbo system for 7 min at 2.5 A and 25 V. Membranes were blocked in 2.5% (w/v) skimmed milk in PBST either overnight at 4 °C or for 1 hour at room temperature, then washed three times for 5 min in PBST. For detection of His-tagged proteins, membranes were incubated for 1 hour at room temperature with HRP-conjugated anti-Penta-His antibody (diluted 1:2000 in PBS) (QIAGEN), followed by PBST washes. For PDI detection, membranes were incubated overnight at 4°C with rabbit anti-PDI antibody (Cell Signalling Technologies) diluted in PBS, washed in PBST, and then incubated for 2 hours at room temperature with IRDye 680RD Goat Anti-Rabbit IgG (diluted 1:10,000 in PBS) (LICORbio™), followed by further PBST washes. Membranes were rinsed in PBS prior to imaging. His-tagged proteins were visualized using SuperSignal™ West Pico PLUS Chemiluminescent Substrate (Thermo Scientific) and the Synergene G: Box gel doc system, while PDI blots were imaged using a fluorescence-based detection system in the Odyssey Fc Imager (LICORbio). Transfer efficiency was assessed by Ponceau S staining following imaging.

### Co-expression of SVMPs with protein disulfide isomerase

The PII and PIII SVMPs were additionally co-expressed with human or snake protein disulfide isomerase (PDI) which was incorporated into the plasmid using Cre-lox (*67*). The multigene expression construct was made using PDI-containing pIDC plasmids (Geneva Biotech). pIDC_hPDI and pIDC_sPDI (PDI-insert synthetised by Genscript) as the donor plasmids were fused with SVMP-containing pACEBac1 plasmids as the acceptor plasmid (*67*). SVMP expression was carried out as above.

### Recombinant SVMP purification

Clarified media (containing SVMP) was continuously passed through HiTrap IMAC FF 5 ml column (Cytiva) at 2 ml/min, for 18 hours at 4°C using a peristaltic pump. Subsequently, the column was washed with 10 CV High Salt Buffer (50 mM Tris-Cl, 1 M NaCl, pH 8.0) and 10 CV SVMP Buffer (50 mM Tris-Cl, 300 mM NaCl, pH 8.0). After the washing steps, the IMAC column was attached to an AKTA Pure system. The SVMP was eluted from the IMAC column using a gradient of imidazole from 0 mM to 500 mM imidazole in SVMP buffer. Fractions containing SVMP were pooled and dialyzed into SVMP Low Salt Buffer (50 mM Tris-Cl, 50 mM NaCl, pH 8.0). SVMP was further purified by ion exchange chromatography by passing the dialyzed protein through a 5 ml HiTrap Q XL column. Subsequently, the SVMP was eluted from the column by a gradient of 50 mM to 1 M NaCl in SVMP buffer. Fractions containing SVMP were pooled and concentrated to 500 μl (maximum 10 mg/ml). Finally, SVMP was purified by size exclusion chromatography (SEC) using an Superdex 200 increase 10/300 GL column (Cytiva) in SVMP buffer with a flow rate of 0.375 ml/min. Cytiva Gel Filtration Calibration Kits were used to calibrate the column, with elution peaks plotted to generate a standard curve with which to calculate the molecular weights (MWs) of SVMP proteins. Eluted SVMP was concentrated to 0.5 - 2 mg/ml using an Amicon concentrator with appropriate MW cut off. Glycerol was added to 5% before the protein was flash frozen in liquid nitrogen and stored at -80°C.

### SVMP deglycosylation assay, PNGase F treatment

Glycosylation of SVMP zymogens was determined by incubation with peptide-*N*-glycosidase F (PNGase F, Promega) to remove N-linked oligosaccharide groups, followed by SDS-PAGE. Briefly, the reaction was carried out with 1-2 μg SVMP diluted in 12 μl water, followed by addition of 1 μl 5% SDS and 1 μl 1 M DTT. Next, the sample was heated at 95°C for 5 minutes to denature the SVMP zymogens. Once cooled at room temperature, 2 μl SVMP Buffer, 2 μl 10% NP-40 and 2 μl PNGase F were added. The reaction mix was incubated at 37°C for 3 hours. Subsequently, SDS-PAGE sample buffer (reducing) was added, and the samples were analysed by SDS-PAGE. SVMP zymogen deglycosylation was identified by a shift towards lower MW of the SVMP band compared to the band where PNGase F was omitted.

### Purification of PIII SVMP fraction from *Echis ocellatus* venom

As a procoagulant control for blood clot time assays, an *Echis ocellatus* SVMP PIII fraction was prepared using SEC. Whole venom (25 mg sourced from the herpetarium facility at the Liverpool School of Tropical Medicine) was dissolved in 1.5 ml of PBS (25 mM sodium phosphate, 150 mM NaCl, pH 7.2), centrifuged and the supernatant loaded onto a 120 ml column of Superdex75 (Cytiva). Proteins were separated in PBS using a flow-rate of 1.0 ml/min and the separation was monitored at 280 nm. Two ml fractions were collected and SDS-PAGE was used to determine the fractions containing SVMP PIIIs. These were pooled and shown to be free of serine protease activity using the casein assay (see above) with appropriate inhibitors.

### SVMP enzyme activation

To determine optimal activation conditions, SVMP was titrated with 0 – 750 μM ZnCl_2_, then incubated at 26°C or 37°C for 18 hours. Activation was monitored using SDS-PAGE and disappearance of the SVMP zymogen band in Coomassie-stained gels.

### Casein and fibrinogen degradation assays

Activated SVMP was titrated into a freshly prepared 20 μl reaction mix containing casein or fibrinogen at a final concentration of 0.525 mg/ml in SVMP Buffer. Final SVMP concentrations ranged from 0 to 2.34 μM (based on SVMP zymogen concentration at the start of the experiment). The negative control contained 5 mM EDTA as an SVMP inhibitor and maximum concentrations of SVMP (PI - 2.34 μM, PII -1.07 and 1.34 μM, PIII - 685 nM). Incubation was carried out at 37°C for 18 hours. SDS-PAGE sample buffer was added, and the samples were heated to 95°C to stop the reaction, and the degradation pattern was analysed by reducing SDS-PAGE.

### Fluorogenic peptide substrate cleavage assay

The quenched fluorogenic peptide Mca-KPLGL-Dpa-AR-NH_2_ (ES010, R&D systems Inc.) was used to kinetically measure SVMP activity, whereby the activated SVMP cleaves the Gly-Leu bond. Cleavage leads to the release of the Dpa-containing quencher from the fluorescent Mca-containing fragment. ES010 substrate was diluted in reaction buffer (50 mM Tris-Cl, 300 mM NaCl, pH 7.5) and used in the assay at final concentrations of 0 μM to 180 μM, in 100 μl in a 96 well plate. Enzymes were activated with Zn^2+^ (see above) and added to a final concentration of 40 or 20 nM based on the SVMP zymogen concentration at the start of the experiment. Fluorescence was followed for 30 minutes or 1 hour at 25°C using a BioTech Synergy Neo2 instrument, an excitation wavelength of 320 nm and emission wavelength of 405 nm. Measurements were taken in triplicate. Kinetic data was plotted using Origin (RRID:SCR_014212, https://www.originlab.com/index.aspx?go=PRODUCTS/Origin), and a non-linear curve was fitted using the Michaelis-Menten function. Origin provided the best-fit values for V_max_ and K_M_, and k_cat_ could be calculated by V_max_/enzyme concentration. The fluorescence intensity versus time curve and the reaction rate were plotted and calculated using GraphPad Prism 10.

### Insulin B degradation assay

Insulin B degradation was determined following the established protocol (*15*), using a final concentration of 0.02 mg/ml insulin B and 0.01 mg/ml activated SVMP, in 50 mM Tris-Cl, 150 mM NaCl, 1 mM CaCl_2_, pH 7.5 (TBS) buffer. Incubation was carried out at 37°C for 120 minutes, then trifluoracetic acid was added to 1% to stop the reaction. The whole sample (= 1 μg insulin B) was analysed by RP-HPLC using a C18 reverse phase column.

### Plate-based aggregometry

Research involving derivatives of human blood samples was approved by the United Bristol Healthcare NHS Trust Research Ethics Committee (project E5736), and all participants provided written informed consent in accordance with the Declaration of Helsinki. Blood samples were collected from healthy adult volunteers into 3.2% sodium citrate Vacutainer tubes, and platelet-rich plasma (PRP) was prepared by centrifugation of citrated whole blood at 180 × g for 17 minutes at room temperature. 2-MeS-ADP was prepared in HEPES-Tyrode’s buffer (10 mM HEPES pH 7.3, 145 mM NaCl, 1 mM MgSO_4_, 3 mM KCl), and experiments were performed using an end-point assay in 96-well half-area plates. 5 µL of each SVMP (PI, PII, PIII and PII zymogen) were incubated with 45 μL PRP for 5 min at 37 °C prior to platelet aggregation induction by 2-MeS-ADP (2.5 μM). The plate was shaken at 1200 rpm, 37 °C for 5 min, and the absorbance was then read at 595 nm using a Labtech LT4500 plate reader. To determine % aggregation, the sample absorbance was normalized accounting for unstimulated PRP as 0% aggregation, and the control (buffer without SVMP) as 100% of aggregation. All means were calculated based on three assays performed in duplicate. A two-way ANOVA was conducted, followed by simple effects analysis (Dunnett post-test) to compare column means within each row, using GraphPad Prism 10.

### Bovine plasma clotting assay

The assay was carried out following the protocol described by Still and colleagues (*68*). Briefly, 20 μl activated SVMP or EchOce NGA (Nigerian *Echis ocellatus* venom) at 20 μg/ml was added to 40 μl of 20 mM CaCl_2_, and then 40 μl of citrated bovine plasma (Biowest S0260-500) was added immediately before measuring absorbance at 595 nm on a CLARIOstar Plus microplate reader. The clot formation was monitored for 30 minutes, at room temperature. To obtain the normal clotting curve, 20 µl of PBS was added. For the no clotting control, CaCl₂ was replaced with 20 µl of PBS. Absorbance versus time curves were plotted using GraphPad Prism 10.

### Thrombin / SVMP clot time assay

Experiments were carried out following manufacturer’s kit instructions (HemosIL, Thrombin Time 0009758515), using an ACL TOP coagulation analyser using 3.0 U/ml thrombin, 500 ng PIII SVMP pool purified from *Echis ocellatus* venom and 500 ng activated recombinant PIII SVMP. The assay measures the rate of increase in turbidity as the fibrin clot forms. The peak in the rate of change (2^nd^ derivative) is considered to be the point of clot formation.

### MTT cell viability assay

MTT assay was based on the methods of Hall *et al.* (*47*). Immortalized human epidermal keratinocytes, HaCaT (Caltag Medsystems, cells derived from a 62-year-old Caucasian male donor’s skin epidermis, RRID: CVCL_0038) were seeded (5,000 cells/well, clear-sided 96-well plates) in standard medium (DMEM high glucose with GlutaMAX^TM^ supplemented with 2 mM sodium pyruvate, 100 IU/ml penicillin, 250 µg/ml streptomycin, 10% Fetal Bovine Serum, 1% GlutaMAX^TM^ (Thermo Fisher Scientific, #35050061)), then left to adhere overnight at 37°C, 5% CO_2_. The cell line was freshly procured and checked for contamination before the start of the experiment. The next day, the SVMPs treatments (25, 50 and 100 µg/ml) were prepared in standard medium, and cells were treated with each prepared solution (100 µl/well, duplicate wells) for 24 hours. For this assay, the MTT Cell Viability Assay Kit (Biotium) was used: 10 μl of MTT solution was added to the 100 μl of medium in each well and mixed by tapping gently on the side of the tray. The cells were incubated at 37 °C for 4 hours. 200 μl DMSO was added directly into the medium in each well and pipetted up and down several times to dissolve the formazan salt. The absorbance was measured on BioTek Synergy Neo2 spectrophotometer at 570 nm. The background absorbance was also measured at 630 nm (subtracted background absorbance from signal absorbance to obtain normalized absorbance values). The % of cell viability for each treatment well was calculated as follows with buffer-treated cells as a control (100% viability):

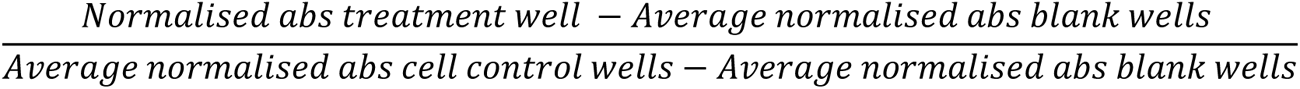

## Materials availability

Materials from this study are available from the corresponding authors upon reasonable request.

## Acknowledgements

The authors thank all members of the Berger, Schaffitzel and Casewell teams for discussions and support. We gratefully acknowledge Loïc Quinton and Thomas Crasset (University of Liège, Belgium) for their expert advice and insightful discussions on mass spectrometry analyses of SVMPs.

## ADDITIONAL INFORMATION

### Funding

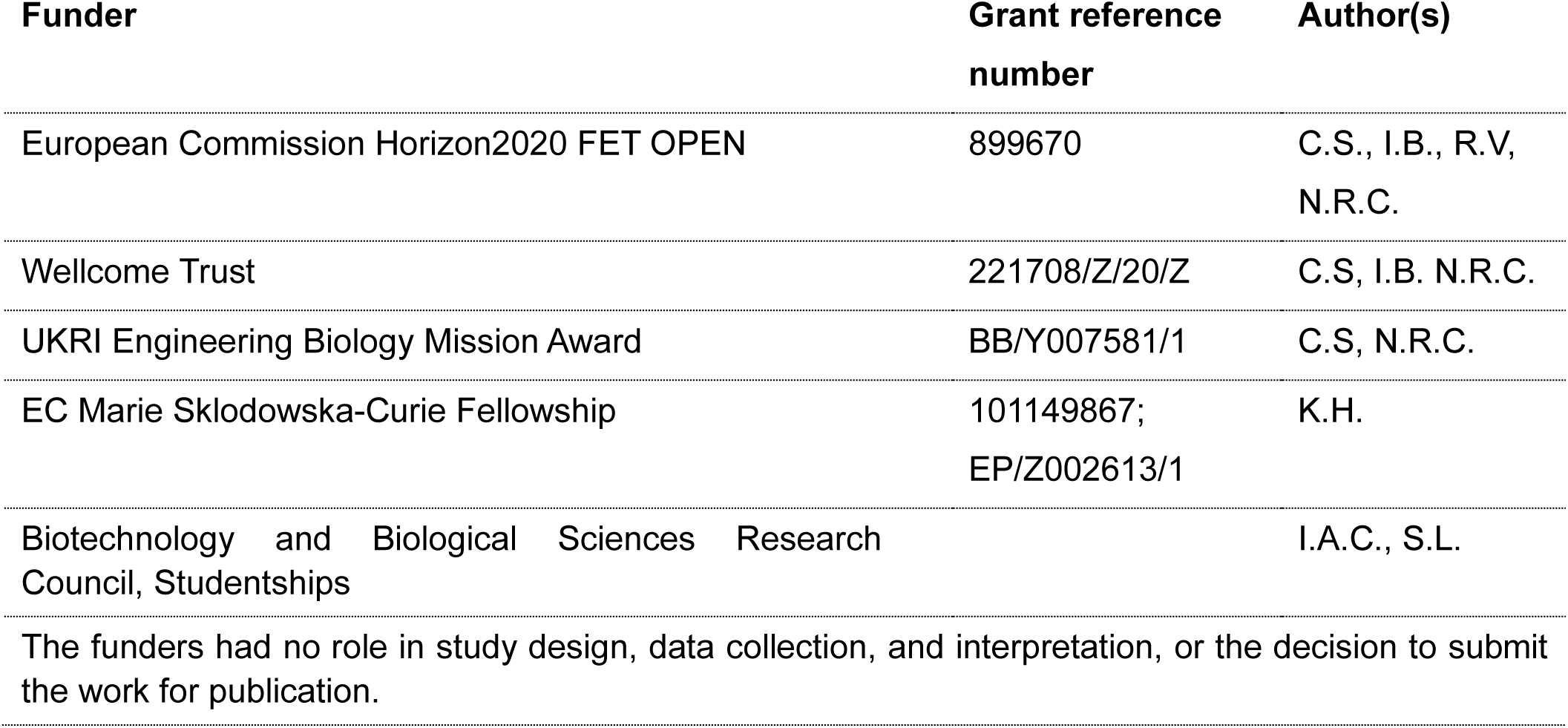

### Author contributions

Sophie Hall, methodology, data curation, formal analysis, investigation, writing – original draft; Iara Aimê Cardoso, methodology, data curation, formal analysis; Mark C. Wilkinson, methodology, data curation, investigation formal analysis, writing - original draft; Maria Molina Carretero, Srikanth Lingappa, Bronwyn Rand, Dakang Shen, data curation, investigation; Johara Boldrini-França, methodology, writing – review and editing; Richard Stenner, methodology; Stefanie K. Menzies, methodology; Georgia Balchin, Konrad Kamil Hus, investigation; Renaud Vincentelli, investigation; Andrew Mumford, Alastair Poole, supervision, writing - original draft; Nicholas R. Casewell, Supervision, writing - original draft; Imre Berger, conceptualization, methodology, supervision, writing - original draft; Christiane Schaffitzel, conceptualization, supervision, writing - original draft.

## ADDITIONAL INFORMATION

### Supplementary files

The supplementary information contains supplementary text and 11 additional supplementary figures.

### Data availability

All data generated or analysed during this study are included in the manuscript and supporting files.

