## Supplementary Text and Figures for "Purified Zymogens Reveal Mechanisms of Snake Venom Metalloproteinase Auto-Activation"

### Optimisation of SVMP expression constructs

Initially, recombinant production of **mature SVMPs**, without the N-terminally fused prodomain, was attempted (Fig. S2a). Protein expression yields, however, were extremely low, primarily due to SVMP cytotoxicity causing cell death immediately upon baculoviral infection. This was evidenced by very low cell viability, with more than 60% of cells dying within 72 hours of expression, and 100% cell death by 96 hours into the expression for the PI and PIII SVMPs (Fig. S2a). Yellow fluorescent protein (YFP) is used in the MultiBac expression system as an indicator of baculovirus performance and a proxy for heterologous protein production (*1*). Typically, YFP fluorescence is followed during expression and cells/proteins are harvested when YFP production reaches a plateau (*1*). Throughout mature SVMP expression, YFP fluorescence remained very low, consistent with pervasive premature cell death caused by the SVMP toxin resulting in low YFP production, and cell lysis leading to release of YFP into the media (Fig. S2a). Additionally, the decrease in YFP readings observed (days 3-5 after proliferation arrest) during expression could also be due to proteolytic degradation of YFP by the mature, active SVMP.

Western blot analysis detecting the His-tag fused C-terminally to the SVMP, showed that the mature PI and PIII had been successfully expressed within the cells. It also indicated that these SVMPs were soluble because they could be detected not only in the whole cell extract (SNP; combined supernatant/pellet) but also in the cleared lysate (SN; supernatant) (Fig. S2a). However, no SVMP was detected in the media sample in the Western blot, indicating that little or no SVMP had been secreted. This is possibly due to the premature death and lysis of the cells, caused by the toxin, impairing the secretion machinery before noticeable secretion had occurred. Taken together, expression of mature SVMPs detrimentally affects cell integrity, damaging the cells early after baculovirus infection and as soon as protein production initiates, underscoring the toxicity of the SVMPs.

**Marimastat** binds to the active site of the metalloproteinase mimicking the structure of a peptide substrate, inhibiting SVMP activity (*2*). During baculovirus infection, Marimastat was added to the medium at a final concentration of 3 µM and 6 µM, respectively (Fig. S2b). At these concentrations however, Marimastat could not prevent cell death induced by PIII SVMP expression. On the contrary, the cell viability in the presence of 3 µM and 6 µM Marimastat was even further decreased. Additionally, less SVMP was detected in anti-His-tag Western blots in whole cell extract (SNP) and cleared lysate (SN), compared to the expression of mature SVMP without

Marimastat (Fig. S2a). Thus, Marimastat itself appears to be toxic to insect cells at the doses tested and has no positive effect on SVMP expression and recovery.

**SVMP active site mutants** were designed based on a previous report of an inactive PII SVMP purified from *Bothriechis lateralis* venom (*3*). Two active site mutations were identified in the protein rendering it inactive (canonical sequence: HEXXHXXGXXH, inactive sequence: H**D**LGHNL**C**IDH). We inserted these mutations into our PI and PIII SVMPs. We found improved cell viability and detected elevated YFP signal for the PI SVMP active site mutant (Fig. S2c). However, the PIII SVMP active site mutant still showed significant toxicity, reducing YFP levels as well as cell viability. Nonetheless, we successfully purified both PI and PIII SVMP active site mutants from the media using immobilized metal affinity purification (IMAC) via the octa-histidine tags present in our proteins. Based on our observations with the active site mutants, we concluded that inhibition of metalloproteinase activity during SVMP protein production has a positive effect, indicating that expressing inactivated SVMPs could be a viable, although inefficient, approach.

Next, we designed **propeptide-SVMP fusion** proteins using the propeptide responsible for the cysteine switch, PKMCGVT. We fused three propeptide repeats to the N-terminus of the SVMP followed by a TEV protease cleavage site. We hypothesized that this would increase the avidity of the propeptide to the MP active site and efficiently inactivate the enzyme. Cell viability for the propeptide-SVMP fusion expression trials was higher at days 3-5 when compared to the mature SVMPs (Fig. S2a and S2d). However, towards the end of expression less than 25% of cells were left alive (day 5), likely limiting protein secretion. Although YFP fluorescence levels for PI and PIII had slightly increased (Fig. S2d) compared to expression of the mature SVMP (Fig. S2a), YFP expression was still significantly lower as compared to a positive control which expressed an unrelated, non-toxic protein. Western blot analysis confirmed that PI and PIII SVMPs were expressed and soluble (Fig. S2d). However, propeptide-SVMP fusion protein yields in the media remained too low to progress to purification.

**Co-expression of protein disulfide isomerases with SVMP zymogens**

We generated expression cassettes for human PDI or snake PDI alongside our SVMPs and used Cre-LoxP recombination to integrate these cassettes into our SVMP expression constructs (*4*). Successful co-expression of human PDI or snake PDI, alongside our SVMPs, was confirmed by Western Blot and mass spectrometry analysis (Fig. S9a). Contrary to our expectation, however, co-expression of PII and PIII SVMP zymogens with these PDIs did not improve the yield of SVMP zymogens: The SEC chromatograms for PII SVMP zymogens after IMAC and IEX purification expressed in the presence and absence of co-expressed PDI aligned perfectly, with identical peak areas (Fig. S9b). Similarly, the SEC chromatograms for purified PIII SVMP zymogens are virtually identical in the presence or absence of PDIs (Fig. S9c). In conclusion, PDI overexpression in insect cells does not enhance the expression and/or folding of our SVMP zymogens and therefore did not result in a noticeable improvement in yield.

It remained possible, however, that the activity of these SVMPs could have been enhanced by the co-expression of the PDIs resulting in improved protein folding, e.g. through optimized disulfide bond formation. When tested against fibrinogen as a substrate, however, the activities of the different SVMPs remained unchanged by PDI co-expression (Fig. S9d,S9e).

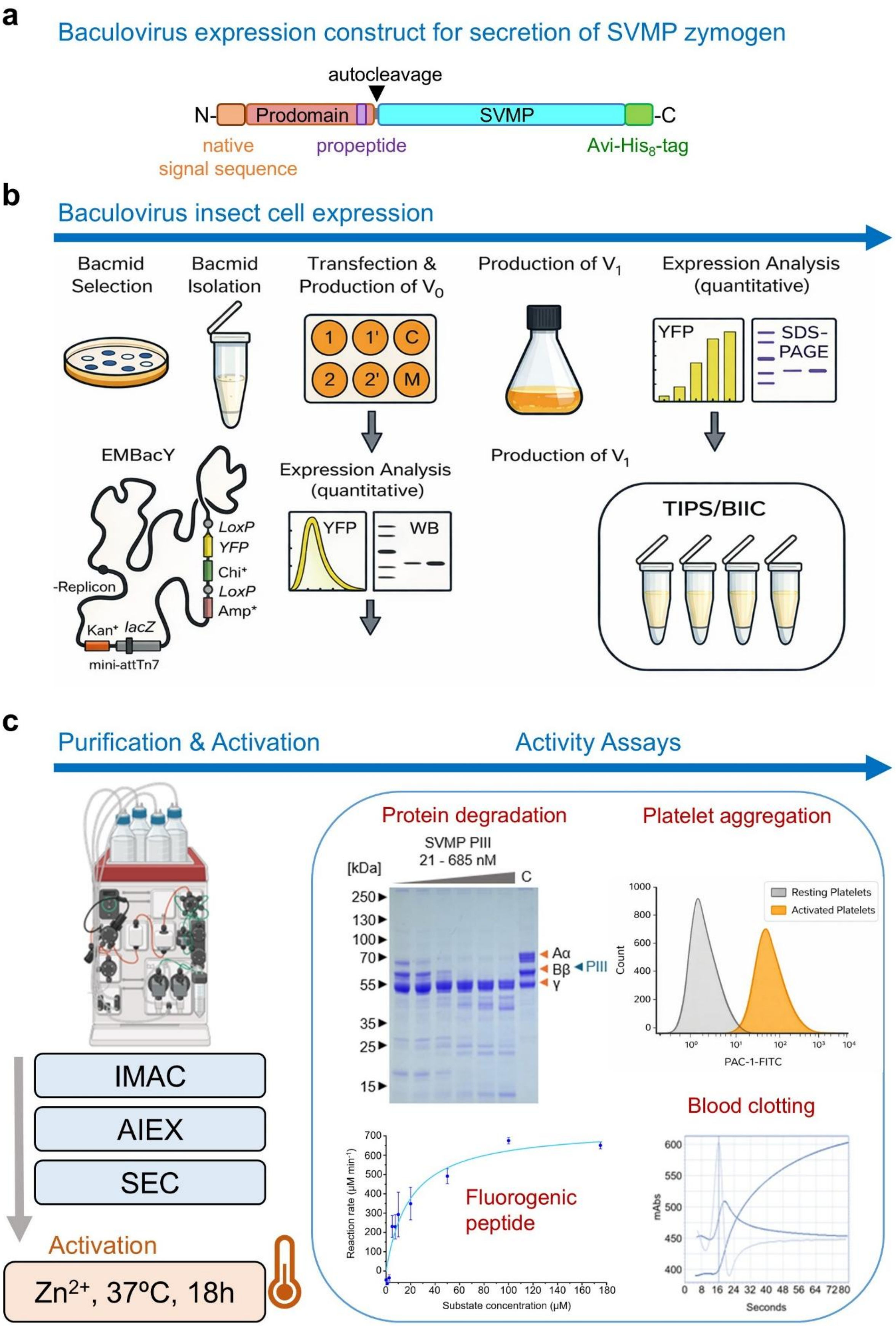
a
Baculovirus expression construct for secretion of SVMP zymogen
autocleavage
N-
Prodomain
SVMP
-C
native
signal sequence
propeptide
Avi-His8-tag
b
Baculovirus insect cell expression
Bacmid Selection
Bacmid Isolation
Transfection & Production of V0
Production of V1
Expression Analysis (quantitative)
1
1'
C
2
2'
M
YFP
SDS-PAGE
EMBacY
LoxP
YFP
Chi+
LoxP
Amp+
-Replicon
Kan+
lacZ
mini-attTn7
Expression Analysis (quantitative)
YFP
WB
Production of V1
TIPS/BIIC
c
Purification & Activation
Activity Assays
IMAC
AIEX
SEC
Activation
Zn2+, 37ºC, 18h
Protein degradation
SVMP PIII
21 - 685 nM
[kDa]
C
250
130
100
70
55
35
25
15
Aα
Bβ
γ
PIII
Platelet aggregation
Resting Platelets
Activated Platelets
Count
PAC-1-FITC
Fluorogenic peptide
Reaction rate (µM min⁻¹)
Substate concentration (µM)
Blood clotting
mAbs
Seconds

**Figure S1: Schematic outlining SVMP production and characterisation strategy.** (**a**) Design of the expression construct for secretion of SVMP zymogens, comprising the native signal sequence, the endogenous prodomain with propeptide, the SVMP and a C-terminal Avi- and octa-histidine tags for biotinylation and affinity purification respectively. (**b**) Baculovirus insect cell expression timeline (adapted from (*5*)). Plasmids encoding the SVMP zymogen constructs are integrated into EMBacY baculoviral DNA via Tn7 transposition. Positive clones are identified by blue/white screening. EMBacY baculoviral DNA (single white colonies) is isolated and used to transfect Sf21 cells. Media containing initial virus ($V_0$) is removed from the wells and used for infecting insect cell cultures (25 ml volume) in an Erlenmeyer shaker flask. Protein production is followed by the YFP fluorescence signal and by Western blot (WB) using anti-His antibodies. Infected Hi5 cell cultures in shaker flasks are split every 24 h until cell proliferation arrest occurs. Media containing amplified virus ($V_1$) is removed ~48 h after pa, and fresh medium is replenished instead. Cells are harvested when the YFP signal has reached a plateau (typically after 3–4 days). Protein production is analyzed by SDS–PAGE. Baculovirus-infected insect cell (BIIC) stocks are prepared for long-term storage of viruses. The whole procedure takes less than 2 weeks. (**c**) SVMP zymogens are secreted and purified from the medium using three consecutive chromatography steps. Activation is accomplished by incubation with $Zn^{2+}$, the concentration of $Zn^{2+}$ required for full activation of the zymogen has to be optimized for each individual SVMP. Activated SVMPs are then used for activity assays, including protein degradation, cleavage of fluorogenic peptide, blood clotting and platelet aggregation assays (for more details see methods). The amount of SVMP required for each assay has to be optimized.

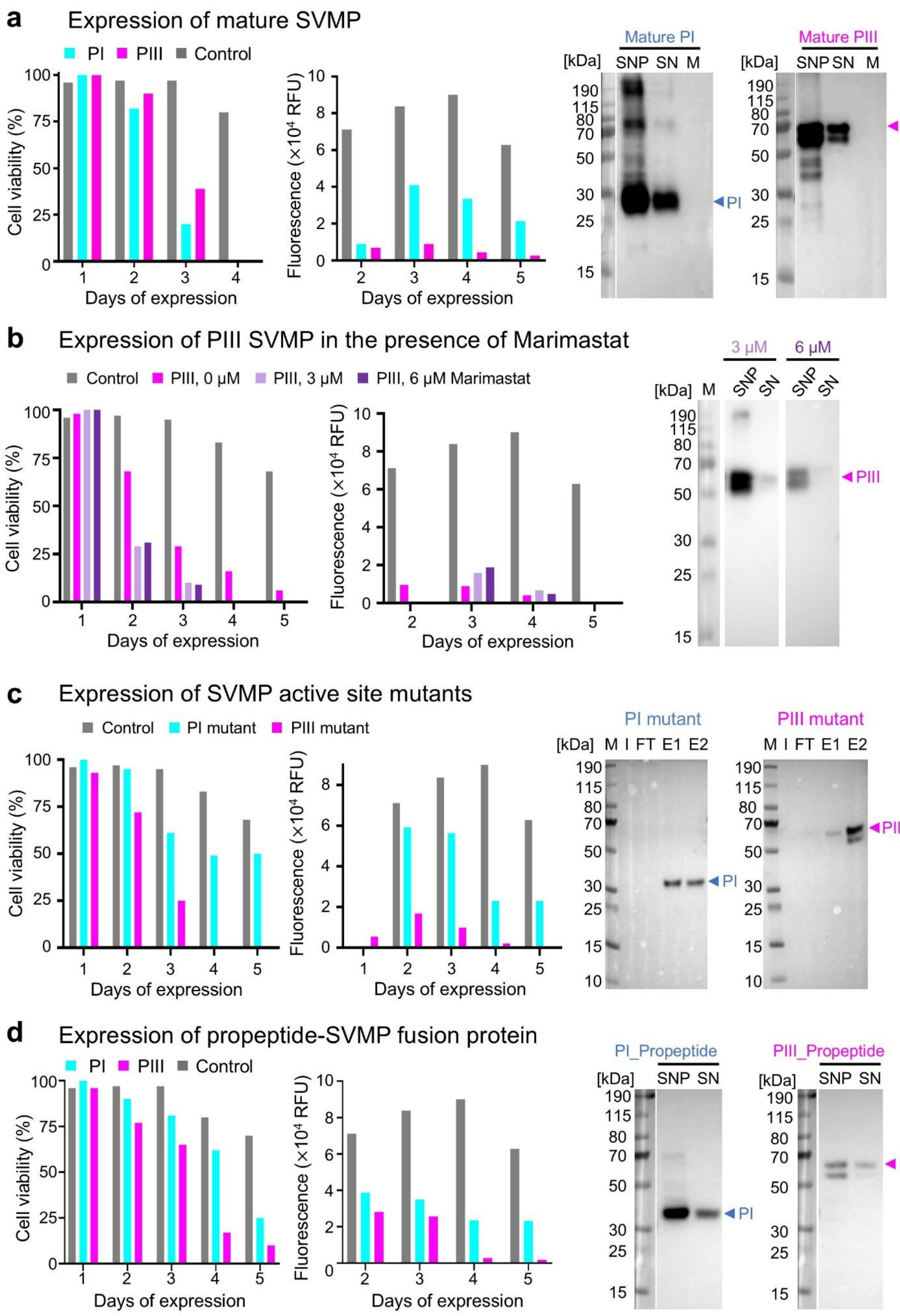
a
Expression of mature SVMP
PI
PIII
Control
Cell viability (%)
Days of expression
Fluorescence (×10⁴ RFU)
Mature PI
Mature PIII
[kDa]
SNP
SN
M
b
Expression of PIII SVMP in the presence of Marimastat
Control
PIII, 0 µM
PIII, 3 µM
PIII, 6 µM Marimastat
3 µM
6 µM
c
Expression of SVMP active site mutants
PI mutant
PIII mutant
M I FT E1 E2
d
Expression of propeptide-SVMP fusion protein
PI_Propeptide
PIII_Propeptide

**Figure S2: Optimisation of SVMP expression** using mature SVMP, broad spectrum inhibitor, SVMP active site mutants and propeptide-SVMP fusion protein to overcome SVMP cytotoxicity. (**a**) Expression of mature SVMPs monitored by cell viability (left), YFP fluorescence (middle), and Western blot analysis of His-tagged protein (right). SNP: supernatant + pellet; SN: supernatant, M: medium. Grey bars: non-toxic protein control expression, cyan: PI SVMP, magenta: PIII SVMP. (**b**) Expression of mature PIII SVMP in the presence of 0 µM (magenta), 3 µM (lilac) or 6 µM (purple) Marimastat. Expression is monitored by cell viability (left), YFP fluorescence (middle), and Western blot analysis of His-tagged protein (right). (**c**) Expression of SVMP active site mutants monitored by cell viability (left) and YFP fluorescence (middle). Right: Western blot analysis of expressed His-tagged protein following Ni-NTA purification. I: input; FT: flowthrough; E1, E2: elution fractions. (**d**) SVMP propeptide-fusion expression monitored by cell viability (left), YFP fluorescence (middle), and Western blot analysis of His-tagged protein (right). Grey bars: non-toxic protein expression control, cyan: PI SVMP, magenta: PIII SVMP. HRP-conjugated anti-Penta-His antibody was used for all Western blots. Experiments were performed at least in duplicate.

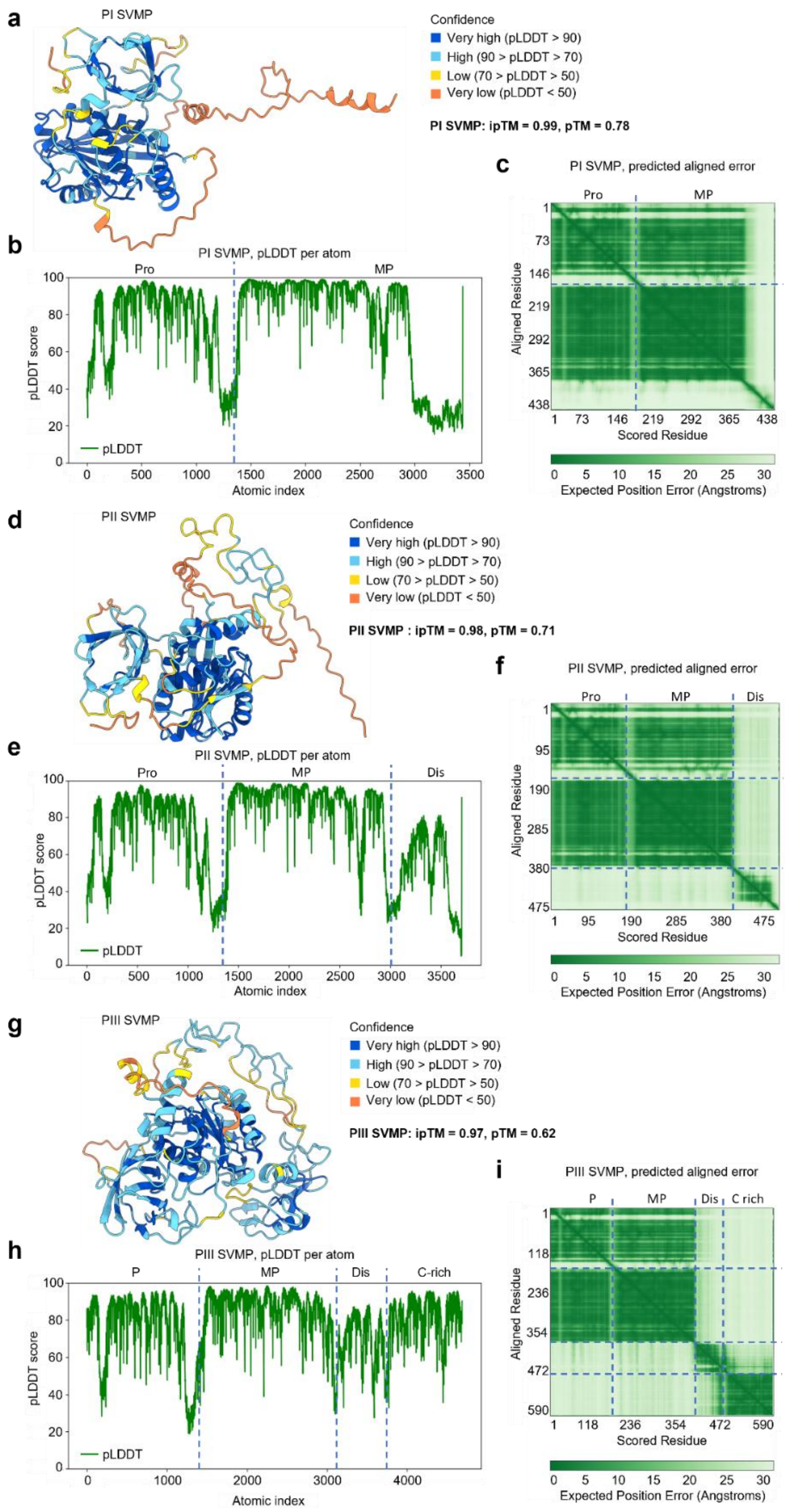


**Figure S3: Structural predictions of SVMP zymogens** of (**a-c**) PI, (**d-f**) PII and (**g-i**) PIII class, performed by AlphaFold 3 (*6*) in the presence of $Zn^{2+}$, along with the corresponding predicted Local Distance Difference Test (pLDDT) (**b, e, h**) and predicted Aligned Error (pAE) (**c, f, i**) plots.

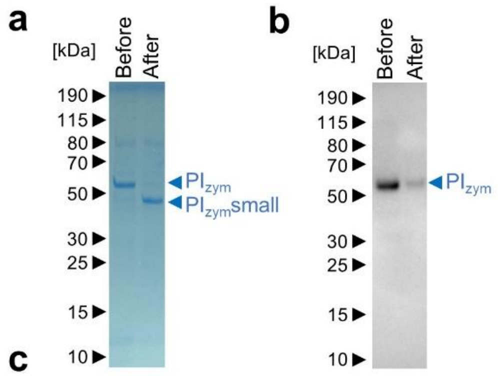


c

| Protein FDR Confidence | Accession | Description | Coverage | # Peptides | # PSMs | # Unique Peptides | Area: F2: Sample | Score Sequest HT |
|---|---|---|---|---|---|---|---|---|
| **High** | **Q2UXQ3+tags** | **PI SVMP** | **59.59** | **27** | **1087** | **27** | **8.532E+10** | **2337.7** |
| High | A0A7E5X2W4 | La-related protein 1-like from *Trichoplusia ni* | 16.22 | 16 | 25 | 16 | 132016395 | 67.07 |
| High | A0A7E5VL90 | Adenylosuccinate lyase from *Trichoplusia ni* | 43.70 | 20 | 23 | 20 | 108915336 | 58.44 |
| High | A0A7E5VKR1 | Vacuolar proton pump subunit B from *Trichoplusia ni* | 25.35 | 11 | 15 | 11 | 53472561 | 36.55 |
| High | A0A7E5WZM0 | ATP synthase subunit alpha from *Trichoplusia ni* | 26.39 | 13 | 18 | 13 | 49168940 | 42.98 |

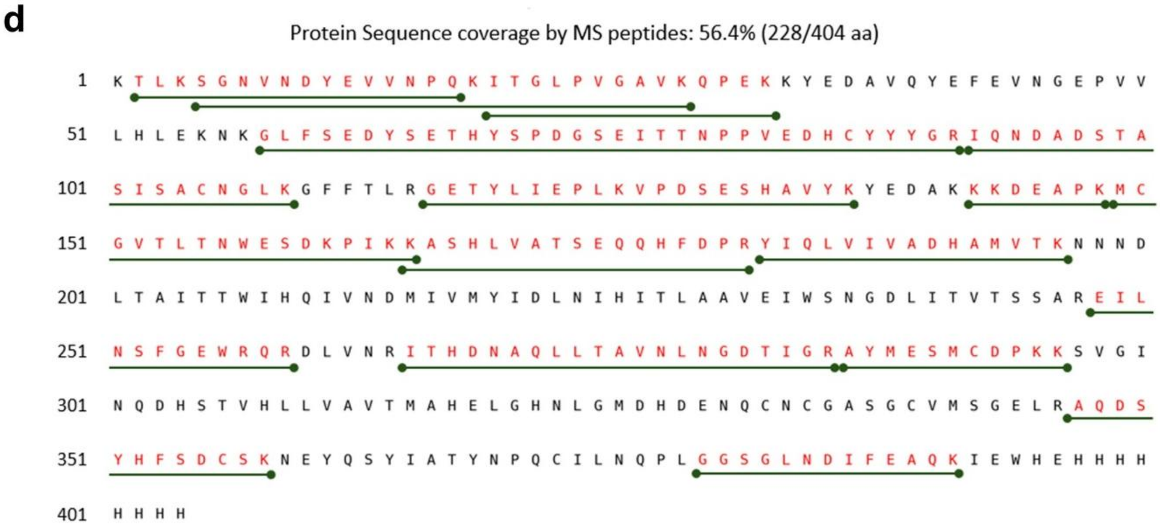


**Figure S4: PI SVMP C-terminal auto-cleavage** before and after dialysis, following IMAC purification. (**a**) SDS-PAGE and (**b**) Western blot using HRP-conjugated anti-Penta-His antibody of sample before and after dialysis, following IMAC purification. (**c**) Top five LC-MS/MS (liquid chromatography-tandem mass spectrometry) hits of before dialysis sample. Proteins were identified using the Sequest search engine against the UniProtKB *Trichoplusia ni* (Hi5) database containing sequence for the protein of interest (PI SVMP). Keratins and trypsin were excluded as common contaminants. (**d**) PI SVMP protein sequence coverage by LC-MS/MS. Peptides identified by mass spectrometry that provide unique sequence information are shown as horizontal bars below the amino acid sequence. Residues detected in the analysis are highlighted in red, covering 56% of the protein sequence.

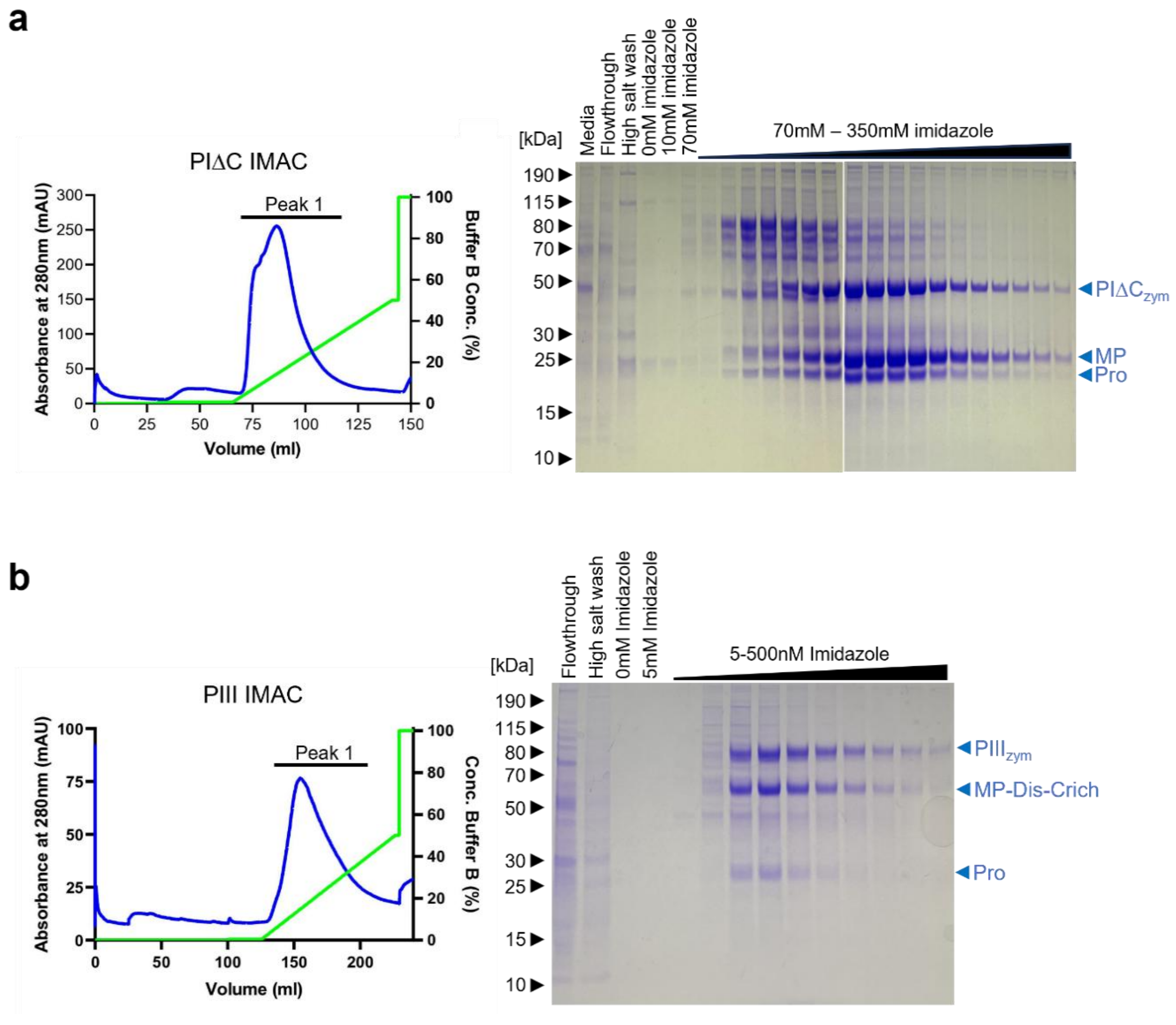


**Figure S5: IMAC purification of secreted SVMPs** from insect cell media. Chromatograms and reducing SDS-PAGE of IMAC purification of **(a)** PIΔC and **(b)** PIII SVMP.

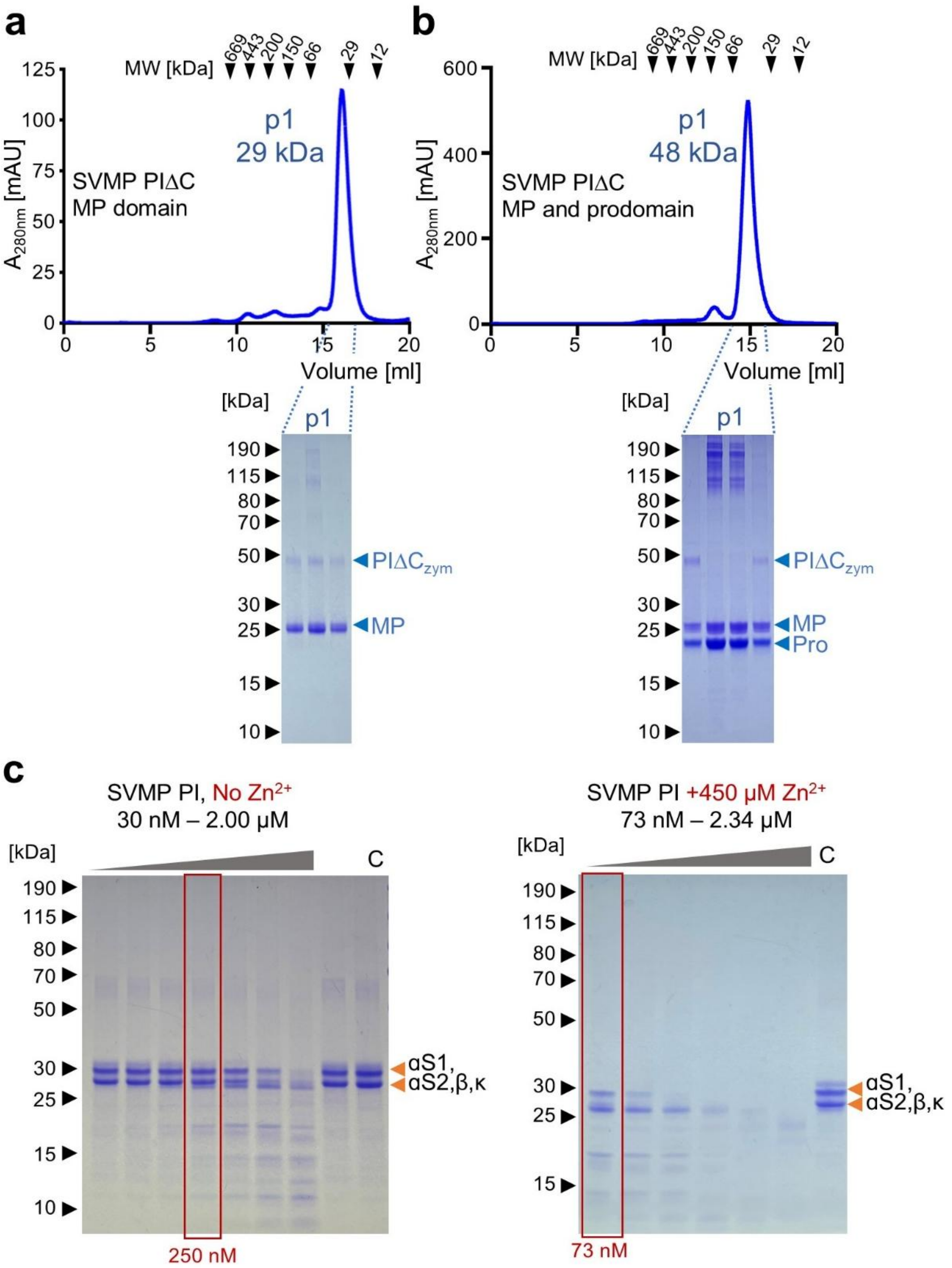


**Figure S6: SEC purification and auto-activation of PIΔC SVMP zymogen,** following IMAC and IEX. Size exclusion chromatograms and SDS-PAGE of (**a**) metalloproteinase domain (MP) of PIΔC and (**b**) metalloproteinase domain (MP) and prodomain (Pro) of PIΔC. (**c**) PIΔC activity against casein with no prior addition of $Zn^{2+}$ ions (left) and with prior incubation with 450 μM of $Zn^{2+}$(right, see also Fig. 4a). Titration of PIΔC SVMP (concentration range indicated) into 23 μM casein, followed by incubation at 37°C for 18 hours. 5μl reaction run on a reducing SDS-PAGE gel.

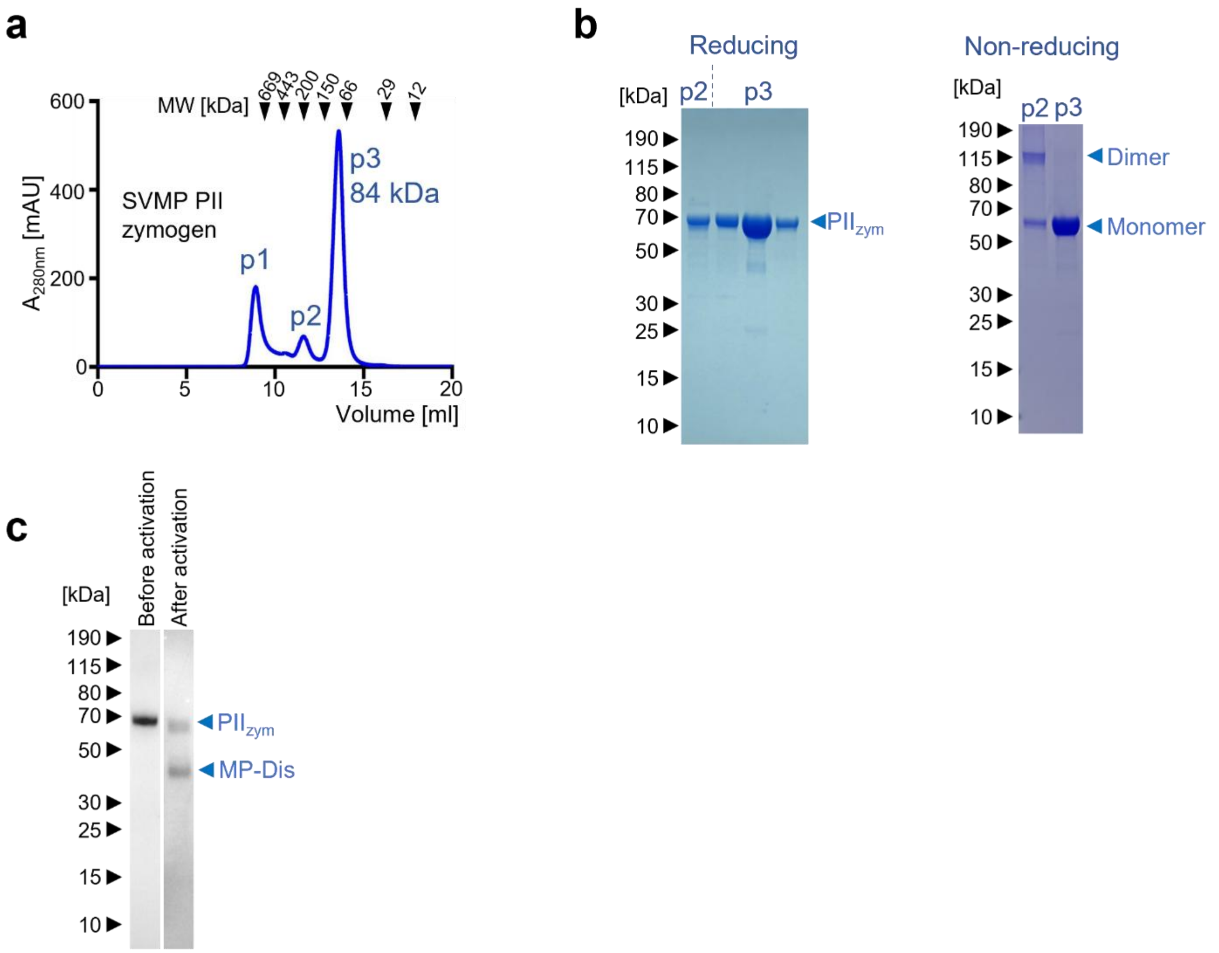


**Figure S7: SEC Purification of PII SVMP zymogen,** following IMAC and IEX. (**a**) Size exclusion chromatogram and (**b**) reducing and non-reducing SDS-PAGE of SEC peaks 2 and 3. (**c**) Western blot analysis using HRP-conjugated anti-Penta-His antibody of sample before and after activation of PII SVMP.

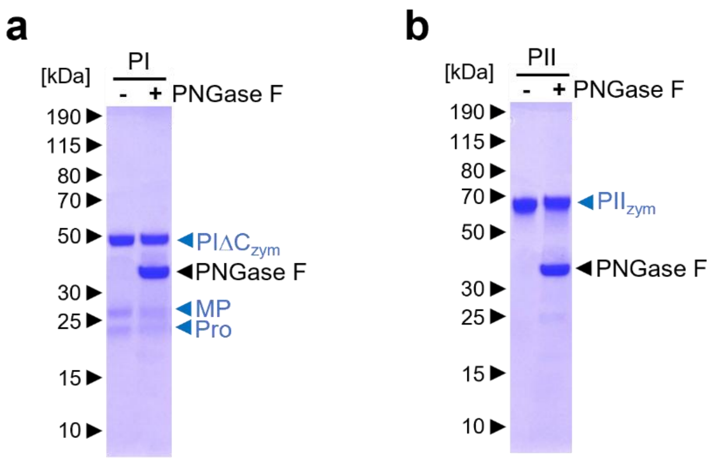


**Figure S8: Glycosylation test of recombinant SVMPs.** Purified (**a**) PI, (**b**) PII and (**c**) PIII SVMP zymogens were treated with PNGase F. A shift to lower MW is observed in bands corresponding to PIII SVMP zymogen and mature PIII SVMP (MP-Dis-Crich) in panel c.

**a**

| Protein FDR Confidence | Accession | Description | Coverage | # Peptides | # PSMs | # Unique Peptides | Area: F4: Sample | Score Sequest HT |
|---|---|---|---|---|---|---|---|---|
| **High** | **sPDI** | **Snake venom protein disulfide isomerase** | **74.76** | **39** | **764** | **39** | **33529221181** | **1699.08** |
| **High** | **PIII SVMP** | **PIII SVMP** | **57.86** | **31** | **108** | **31** | **2499322340** | **308.03** |
| High | A0A7E5VVN0 | T-complex protein 1 subunit delta from *Trichoplusia ni* | 49.91 | 23 | 46 | 23 | 591351696 | 130.49 |
| High | A0A7E5WZQ1 | T-complex protein 1 subunit eta from *Trichoplusia ni* | 54.23 | 24 | 48 | 24 | 560392602 | 125.74 |
| High | A0A7E5X1U5 | T-complex protein 1 subunit beta from *Trichoplusia ni* | 60.82 | 29 | 60 | 29 | 498295832 | 158.19 |

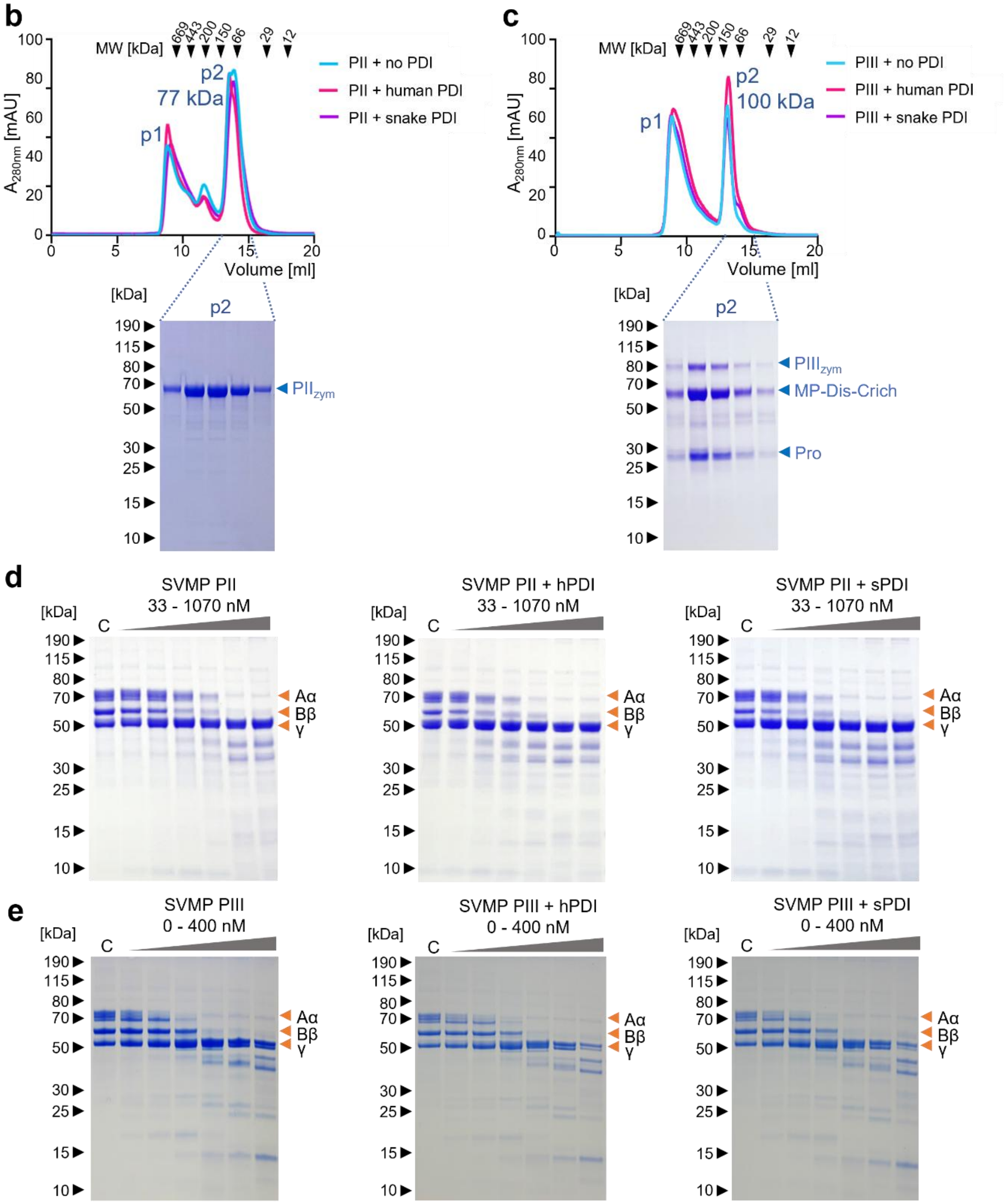

**Figure S9: SEC chromatograms of PII and PIII SVMP zymogens produced in the presence and the absence of human and snake protein disulfide isomerases (PDI).** (**a**) Top five LC-MS/MS hits of PIII + sPDI IMAC flowthrough. Proteins were identified using the Sequest search engine against the UniProtKB *Trichoplusia ni* (Hi5) database containing sequences for the proteins of interest (PIII SVMP and snake PDI). Keratins and trypsin have been excluded as common contaminants. (**b**) PII SVMP zymogen and (**c**) PIII SVMP zymogen were co-expressed with human (magenta) or snake (purple) PDIs. SEC chromatograms were aligned with the control where no PDI was expressed (cyan). (**d**) Fibrinogen degradation assays in the presence of $Zn^{2+}$ and increasing amounts of SVMPs PII, PII + hPDI (human PDI) and PII +s PDI (snake PDI). (**e**) Fibrinogen degradation assays in the presence of $Zn^{2+}$ and increasing amounts of SVMPs PIII, PIII + hPDI and PIII + sPDI. Degradation assays were performed four times for the PII SVMP + PDI assays, and six times for the PIII SVMP + PDI assays.

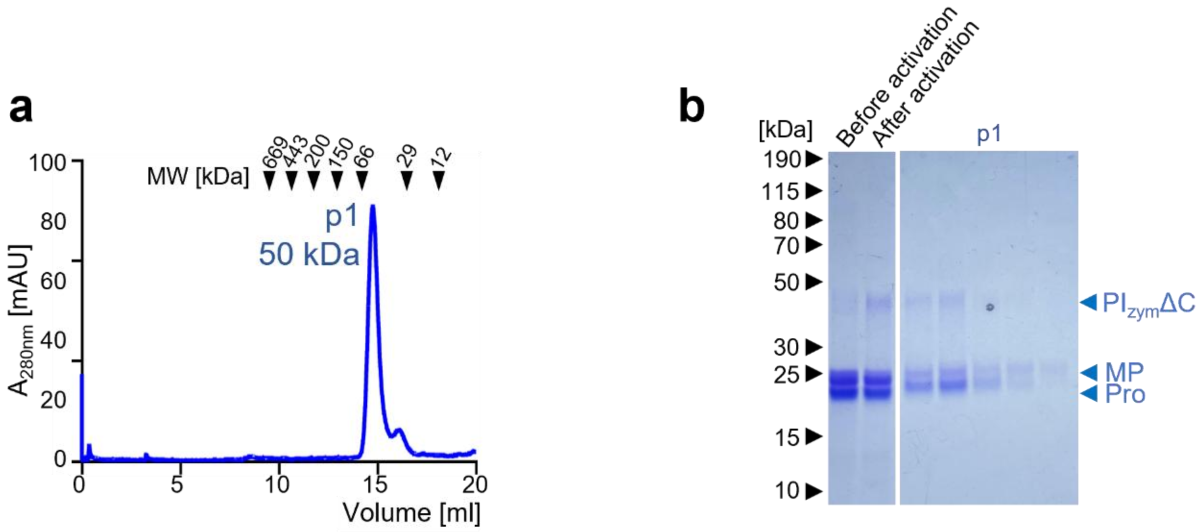


**Figure S10: Auto-activated $PI_{zym}ΔC$ SEC.** (**a**) Size exclusion chromatogram and (**b**) corresponding Coomassie-stained SDS gel of activated $PI_{zym}ΔC$, confirming metalloproteinase domain (MP) and prodomain (Pro) remain bound together, despite the protein being active.

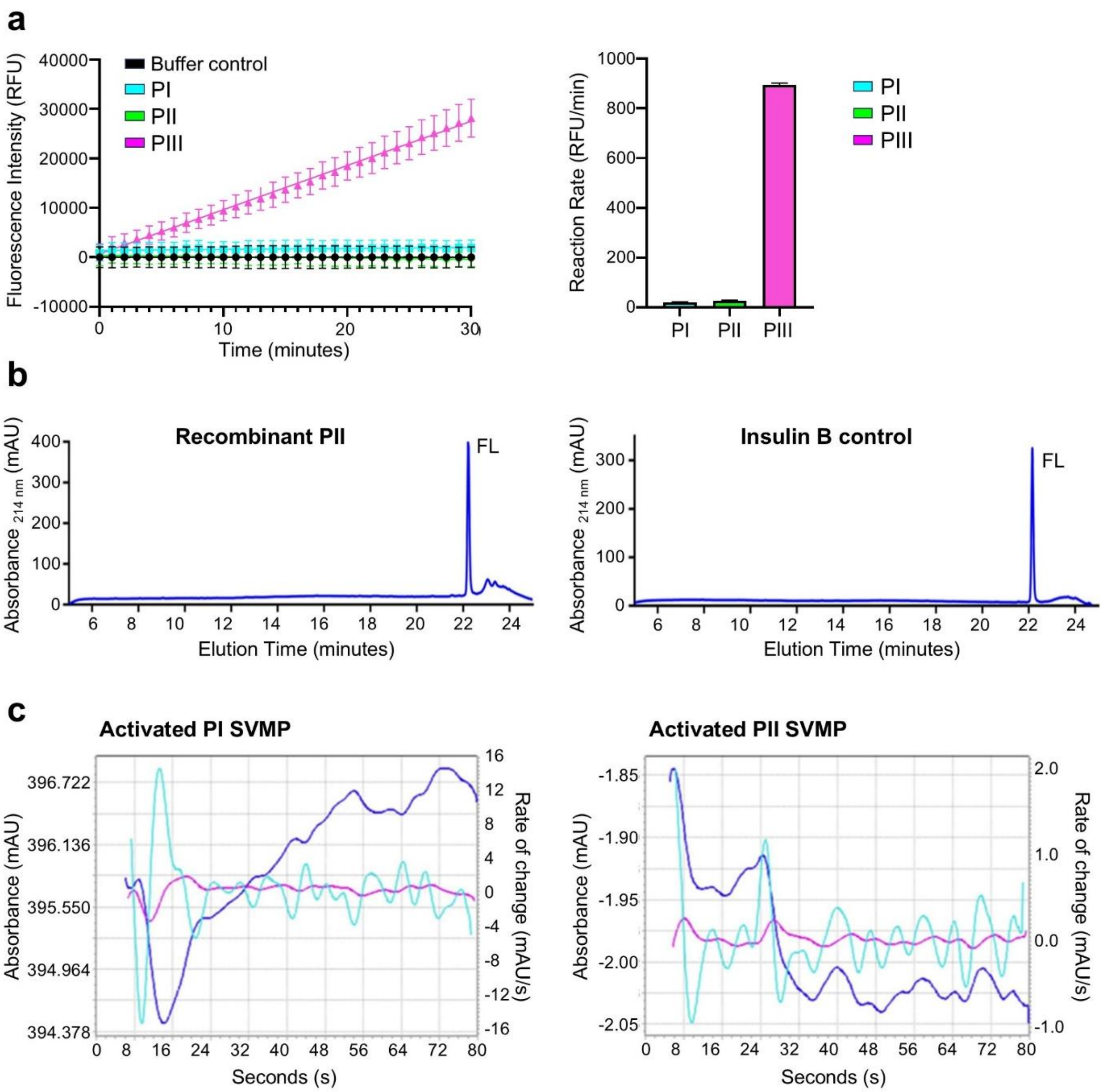


**Figure S11: SVMP activity assays and blood coagulation tests.** (**a**) PI, PII and PIII SVMP activity (40 nM each) towards the fluorogenic peptide substrate ES010 (100 μM). Data points represent the mean of four individual values recorded over two independent technical replicates, and error bars represent standard deviation. The rate of reaction was determined by calculating the slope of the fluorescence intensity versus time curve.(**b**) Degradation of insulin B by PII SVMP (above) and insulin B only (negative control, below). Full-length insulin (FL) is marked. See Figure 4h and 4i PI and PIII SVMP degradation tests, respectively. (**c**) Citrated human blood was spiked with the recombinant PI SVMP (left) and the recombinant PII SVMP (right). Change in absorbance (cyan) is plotted on the left Y axis, rate of change ($1^{st}$ derivative: blue; $2^{nd}$ derivative: pink) is plotted on the right Y axis. See Figure 5d for PIII SVMP for human blood coagulation test.
